# Postmortem Alterations of Metabotropic Glutamate Receptors across Neuropsychiatric Disorders: A Systematic Review

**DOI:** 10.64898/2026.08.29.748003

**Authors:** Ryoma Kani, Tony Kaku, Yu Arai, Takahide Etani, Sunjun Huh, Keisuke Saito, Saki Homma, Koki Takahashi, Masataka Wada, Yasuharu Yamamoto, Nariko Katayama, Takahiro Nemoto, Hiroyuki Uchida, Keisuke Takahata, Shinichiro Nakajima

**Affiliations:** Department of Neuropsychiatry, Keio University School of Medicine, Tokyo, Japan; Saiseikai Utsunomiya Hospital, Tochigi, Japan; Department of Neuropsychiatry, Toho University Faculty of Medicine, Tokyo, Japan; Keio University Hospital, Tokyo, Japan; Department of Psychiatry and Behavioral Sciences, Stanford University, California, U.S; Advanced Neuroimaging Center, Institute for Quantum Medical Science, National Institutes for Quantum and Radiological Science and Technology, Chiba, Japan; Psychiatry Division, Health Center, Keio University, Tokyo, Japan

## Abstract

Metabotropic glutamate receptors (mGluRs) regulate glutamatergic transmission and have been implicated in diverse neuropsychiatric disorders, but human postmortem evidence remains fragmented. We aimed to map these findings across diagnoses, receptor subtypes, brain regions, and measurement modalities. Following PRISMA guidelines, we systematically searched MEDLINE, EMBASE, and Web of Science from inception to August 8, 2026, for studies assessing *GRM* transcripts, as well as mGluR protein abundance, localization, assembly, or receptor binding in human postmortem brain tissue. Of 532 records identified, 57 reports met eligibility criteria. Findings were synthesized narratively because of substantial heterogeneity in diagnoses, brain regions, receptor subtypes, and assays. Postmortem evidence was concentrated on mGluR5, mGluR2/3, and mGluR1, and on the prefrontal cortex, anterior cingulate cortex, and hippocampus. mGluR-related alterations were reported across disorders, including schizophrenia, major depressive disorder, Alzheimer’s disease, autism spectrum disorder, and alcohol use disorder. Although most analyses yielded null findings, the direction and magnitude of mGluR alterations varied across brain regions, receptor subtypes, and molecular endpoints. This inconsistency may partly reflect the distinct biological levels captured by transcript abundance, total protein, receptor assembly, localization, and ligand binding, together with regional, cell-type, disease-stage, and clinical heterogeneity. The available evidence therefore suggests context-dependent alterations in mGluR biology but not a uniform or disorder-specific molecular signature. Integration of postmortem findings with other approaches, including in vivo imaging, may clarify their biological and clinical significance.

## INTRODUCTION

Metabotropic glutamate receptors (mGluRs) are a family of class C G protein-coupled receptors that shape glutamatergic signaling across synapses, cells, and brain circuits. Eight mGluR subtypes encoded by the *GRM* gene family are classified into three groups by sequence homology, signal transduction, and pharmacology: group I (mGluR1 and mGluR5), group II (mGluR2 and mGluR3), and group III (mGluR4, mGluR6, mGluR7, and mGluR8). Unlike ionotropic glutamate receptors, which mediate fast excitatory neurotransmission, mGluRs modulate glutamate release, neuronal excitability, receptor trafficking, intracellular signaling cascades, synaptic plasticity, and excitatory-inhibitory balance [1, 2]. Group I receptors are generally coupled to postsynaptic excitatory signaling and plasticity-related pathways, and group II and group III receptors often regulate presynaptic neurotransmitter release and glutamatergic tone [2, 3], although their localization and function vary across brain regions and cell types [3]. Since glutamatergic dysregulation, synaptic pathology, and circuit-level maladaptation have been implicated in schizophrenia-spectrum disorders, mood disorders, and neurodegenerative disorders, subtype– and region-specific biology is particularly relevant [4, 5].

In vivo imaging and postmortem approaches both address human mGluR abnormalities but measure different aspects of receptor biology. Translation, receptor trafficking, dimerization, and interactions with scaffold proteins may therefore produce differences across measurement levels [2, 6]. In separate cohorts with major depressive disorder (MDD), lower mGluR5 availability measured with positron emission tomography (PET) was accompanied by lower mGluR5 protein abundance in the prefrontal cortex (PFC) [7]. In post-traumatic stress disorder (PTSD), by contrast, higher cortical mGluR5 availability was accompanied in an independent postmortem cohort by unchanged *GRM5* expression but increased *SHANK1* expression, consistent with altered stabilization of mGluR5 at the cell surface [8]. Thus, concordance across assays can strengthen evidence for an mGluR alteration, whereas discordance may reflect altered receptor organization or trafficking rather than a change in total receptor abundance.

Postmortem tissue uniquely enables direct assessment of mGluR transcripts, protein abundance, receptor assembly, cellular localization, and ligand-binding sites that cannot be assessed comprehensively in living participants [9]. Postmortem mGluR evidence, however, remains dispersed across disorders, receptor subtypes, brain regions, and assay platforms, and some publications draw on the same or overlapping donor cohorts, complicating assessment of independent replication and cross-diagnostic patterns. We therefore systematically reviewed human postmortem studies that directly measured *GRM1* through *GRM8* transcripts or mGluR1 through mGluR8 protein expression, localization, assembly, or receptor binding across neuropsychiatric disorders. We aimed to map findings by diagnosis, receptor subtype, and brain region, to explore the relevance of mGluR alterations to disease pathophysiology and mGluR-targeted treatments.

## METHODS

The full protocol was registered on the International Prospective Register of Systematic Reviews website (CRD420261423542). This study followed the Preferred Reporting Items for Systematic Reviews and Meta-Analyses (PRISMA) guidelines (**Supplemental Table S1**) [10].

### Eligibility criteria

The inclusion criteria were the studies [i] utilizing human postmortem brain tissues, [ii] focusing on individuals diagnosed with neuropsychiatric disorders, [iii] investigating mGluR expression, assembly, receptor binding, or mGluR-related transcripts. The exclusion criteria were the studies [i] using non-human animal models or tissues, [ii] measuring the concentrations of substances acting on mGluRs (e.g., agonists, antagonists, or allosteric modulators), [iii] targeting downstream signaling molecules of the mGluR cascade, [iv] being published as reviews, methodological studies, or conference abstracts.

### Information sources and data collection process

MEDLINE, EMBASE, and Web of Science were searched from database inception to August 8th, 2026. The search strategy is shown in **Supplemental Tables S2-4.**

Six investigators (RK, YA, KS, SH, TK, and TE) independently extracted studies meeting the inclusion criteria using Rayyan and assessed the risk of bias using the Newcastle-Ottawa Scale to evaluate case-control and cohort studies [11, 12]. The following information regarding patient and tissue characteristics was collected: age, sex, number of patients, diagnosis, diagnostic criteria, medication exposure, postmortem interval, freezer storage interval, RNA integrity number, and tissue pH. We also extracted data on study characteristics, including assessment methods for GRM transcripts, mGluR expression or receptor binding, target regions, target molecules, and the differences in mGluR-related measures between patients and controls. Two investigators (RK and TK) verified the literature search, data transfer accuracy, and risk-of-bias assessments. Any discrepancies were resolved through discussion and consultation with the corresponding author (SN).

## RESULTS

### Study selection and characteristics

The database search identified a total of 532 articles. After removal of 180 duplicates, 352 articles underwent title and abstract screening, and 212 articles were excluded. Full texts were assessed for 140 reports, of which 83 reports were excluded. The exclusion reasons are shown in the PRISMA flowchart (**Figure 1**). Ultimately, 57 reports were included.

**Figure 1.**
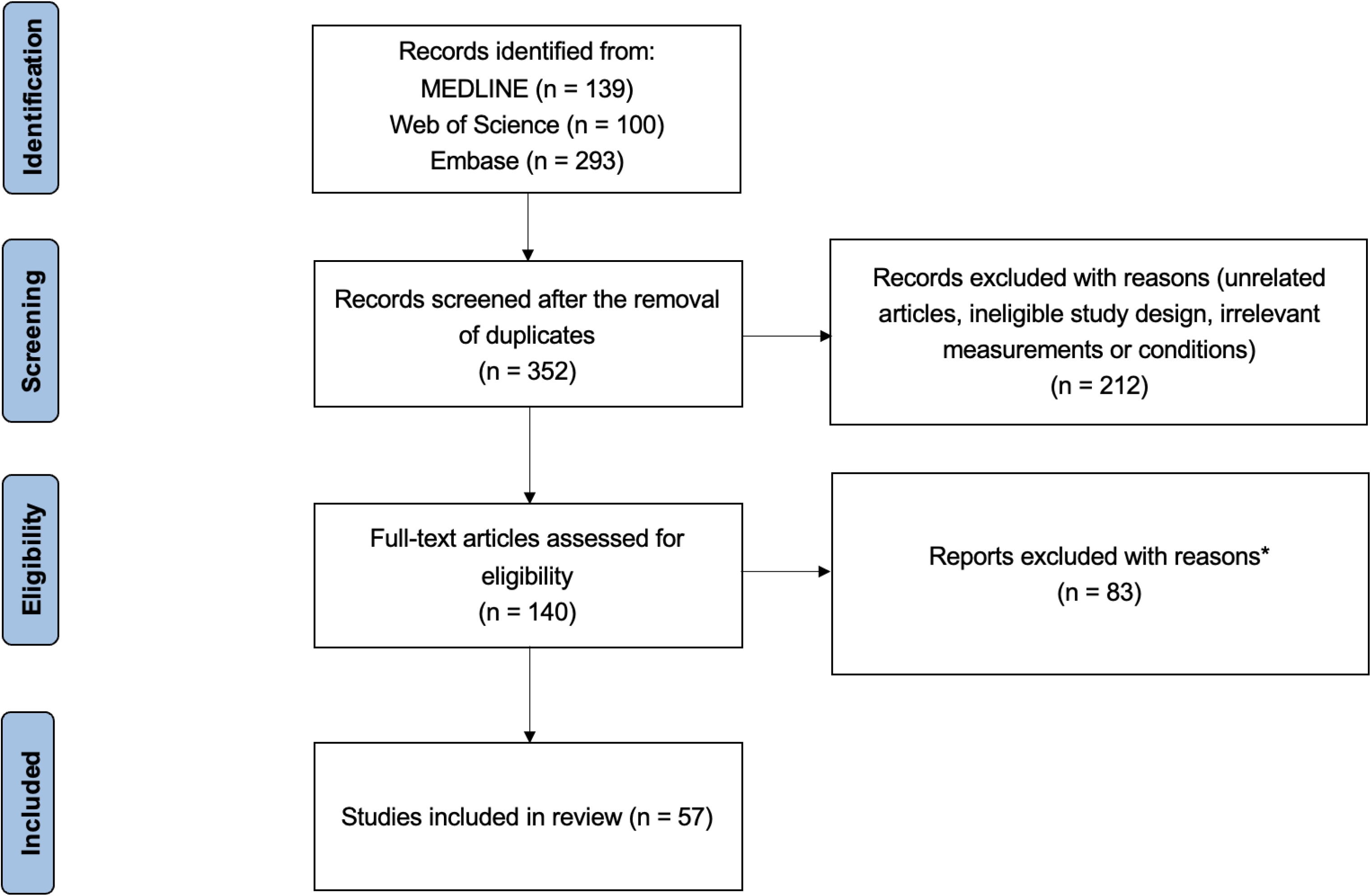
Flowchart of study selection. *Studies because they did not use human postmortem brain tissue (n = 2), did not include eligible neuropsychiatric disorders (n = 20), did not directly assess GRM or mGluR expression, receptor binding, abundance, or localization (n = 15), or did not report mGluR-related measures (n=1), or were reviews, conference abstracts, editorials, or methodological reports (n = 44), or were retracted (n = 1)

The diagnoses, brain regions, methodologies, and medication exposure for each study are detailed in **Table 1**. Distribution of studies by receptor subtype, brain region, and disease group is shown in **Figure 2**. Studies that assessed multiple measurement modalities and/or molecular levels are summarized in **Table 2**. Findings were synthesized narratively rather than pooled quantitatively because studies varied in diagnosis, brain region, receptor subtype, assay platform, and reporting of postmortem variables. Risk-of-bias assessments of included studies were summarized in **Figure S1,** using the NOS-TLPlot [13].

**Figure 2.**
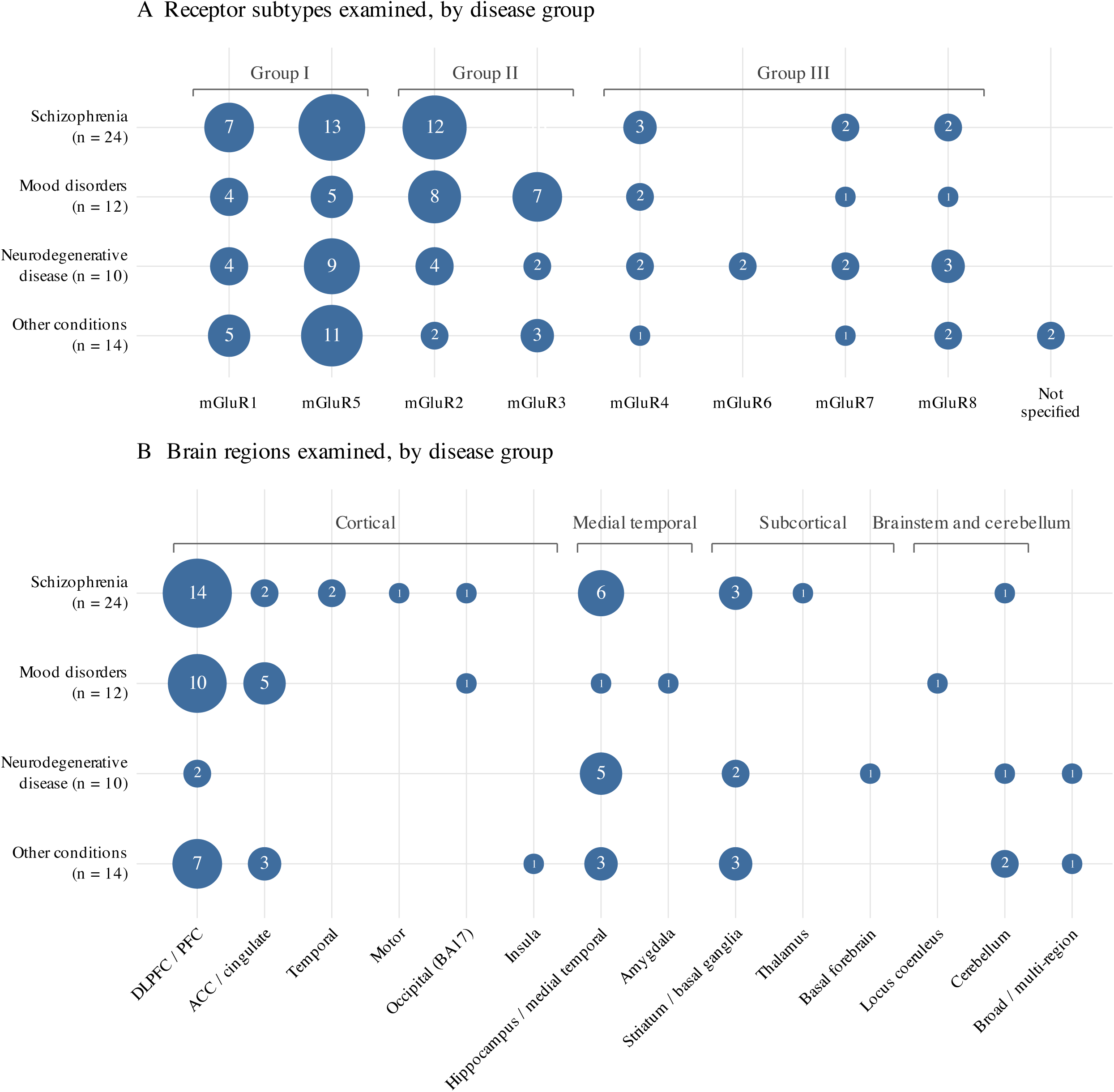
Distribution of studies by receptor subtype, brain region, and disease group. Bubble size represents the number of studies examining each receptor subtype or brain region within each disease group. Abbreviation: ACC: Anterior Cingulate Cortex; BA: Brodmann Area; DLPFC: Dorsolateral Prefrontal Cortex; mGuR: Metabotropic Glutamate Receptor; PFC: Prefrontal Cortex.

**Table 1.**
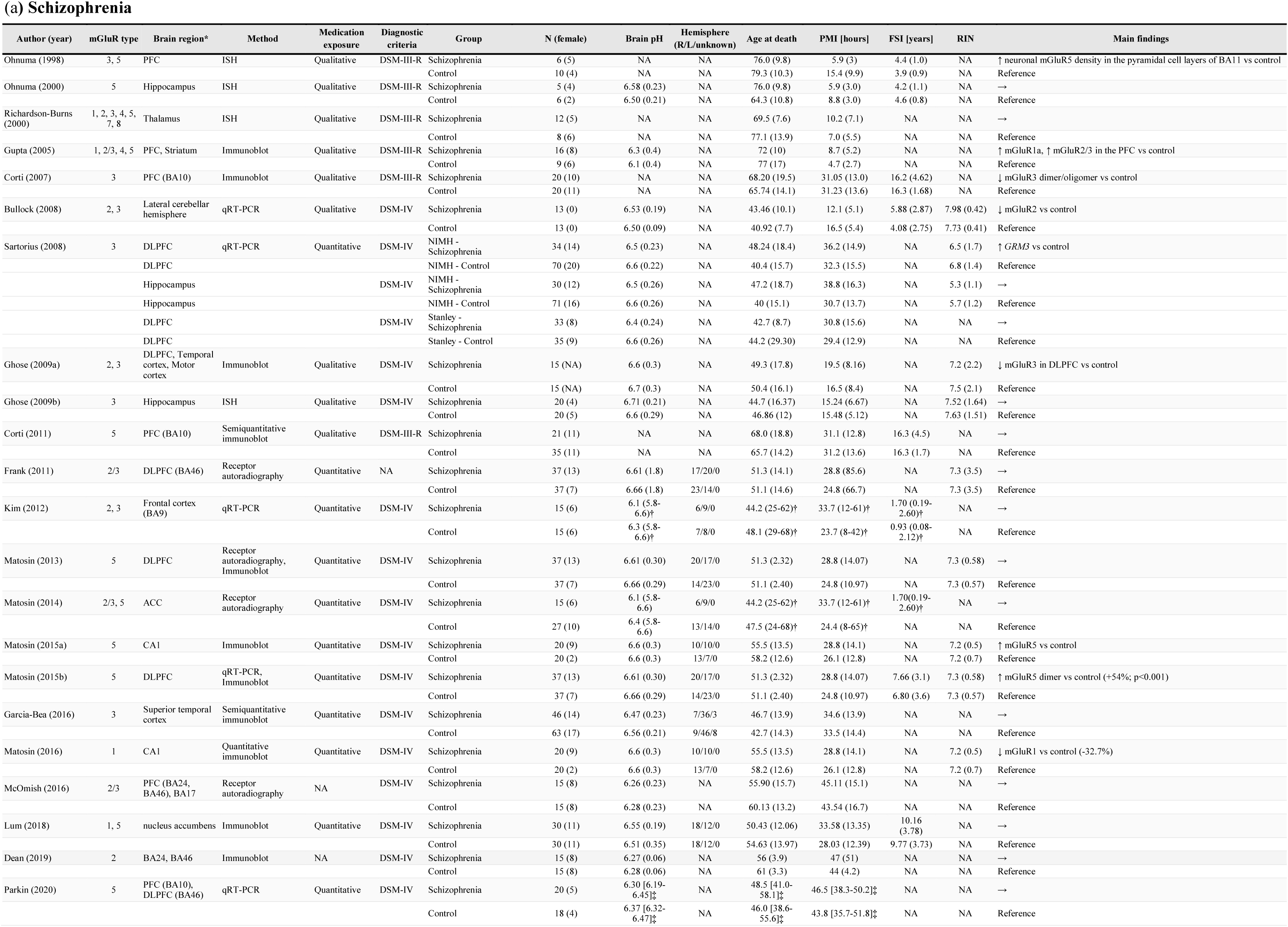
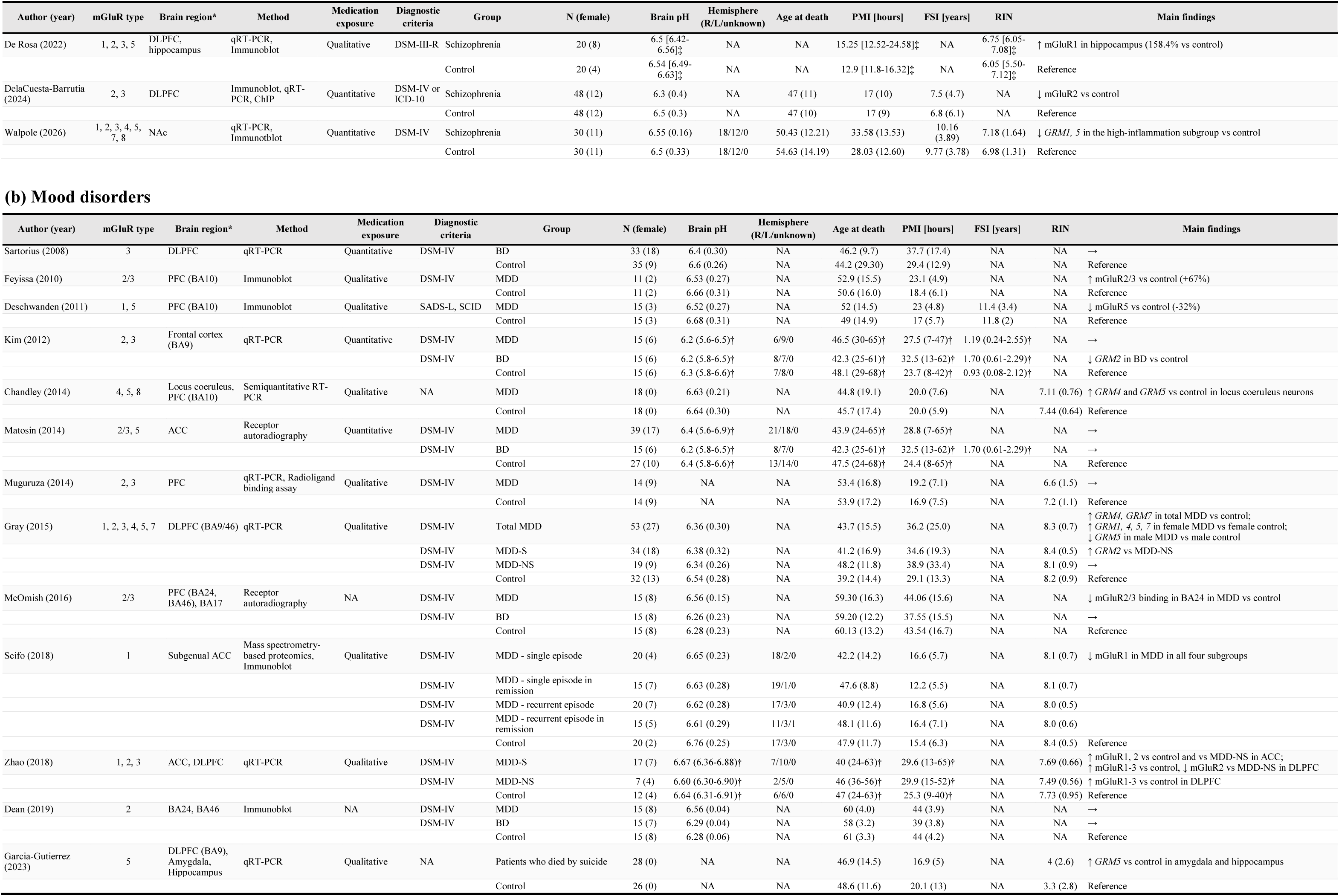
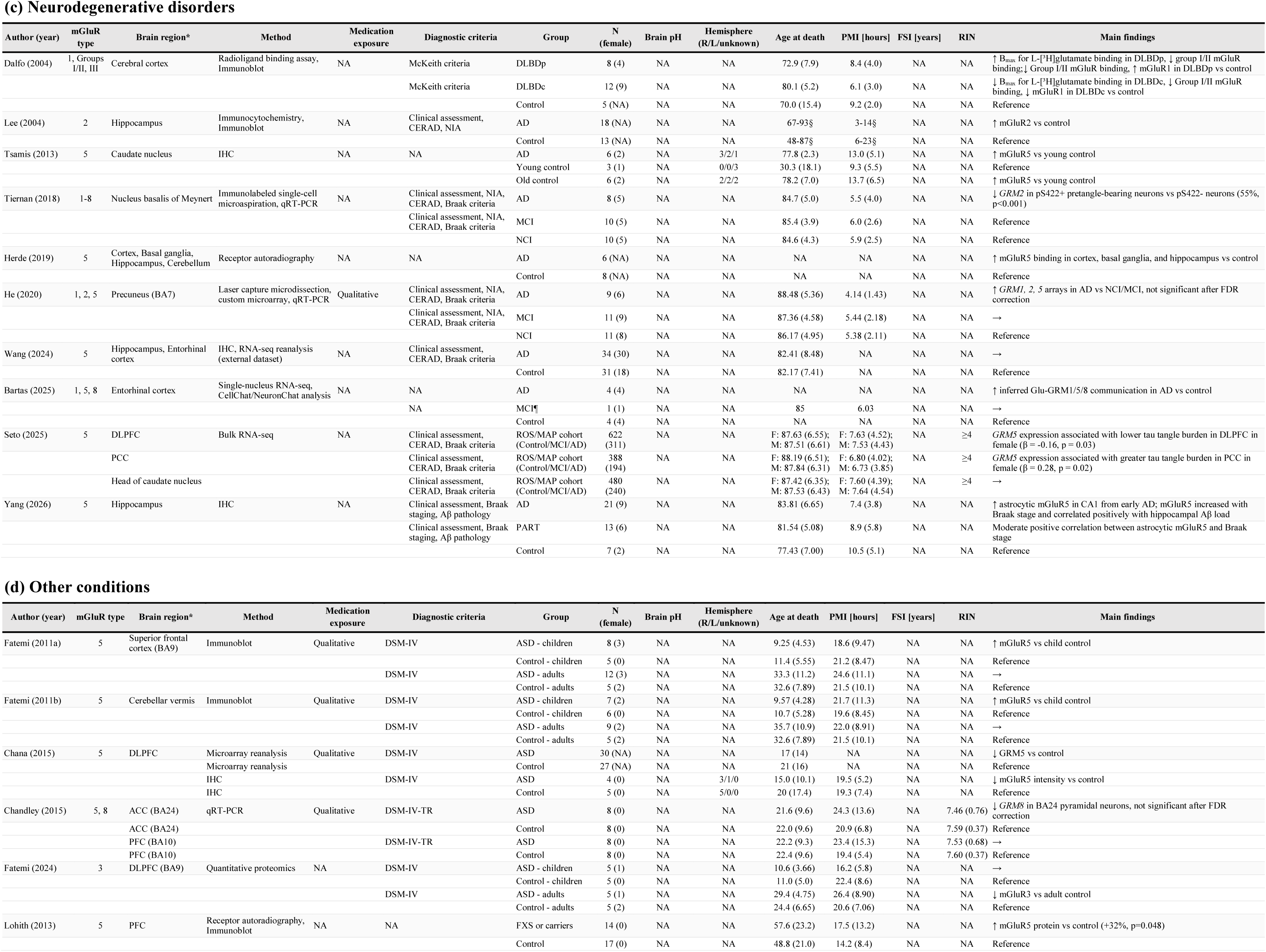
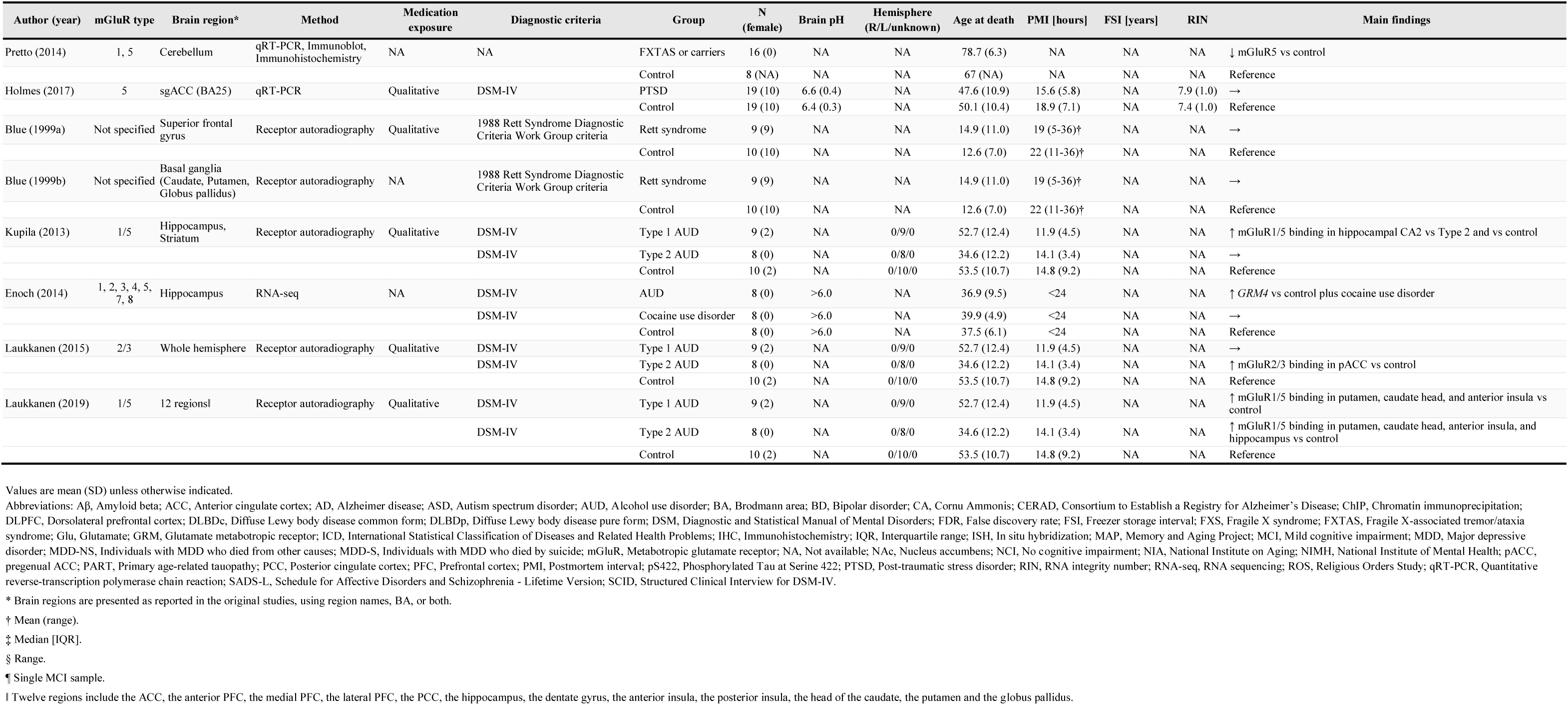
Characteristics of included postmortem studies examining metabotropic glutamate receptors.

**Table 2.**
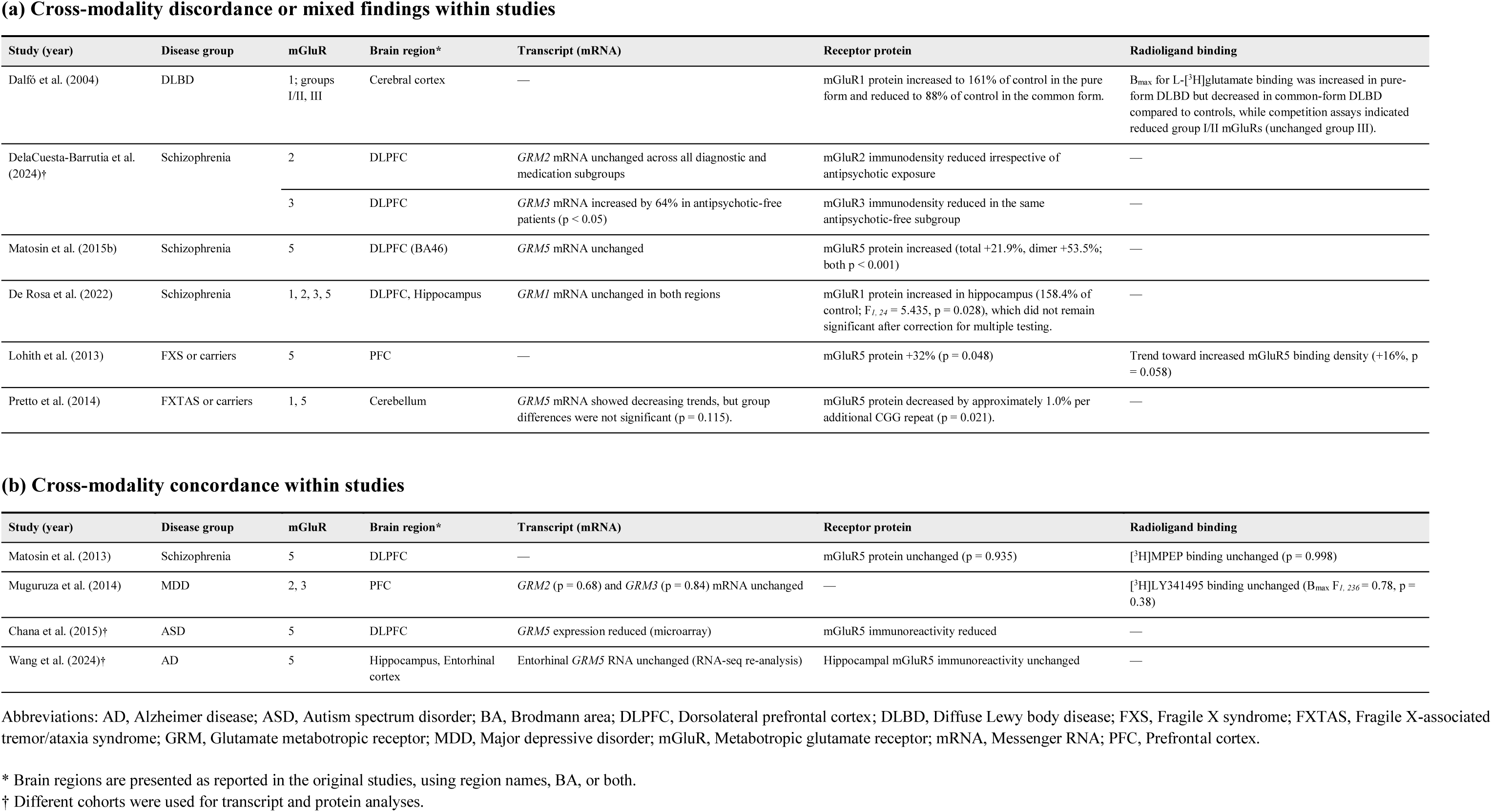
Concordance of mGluR findings across modalities within individual studies.

### Schizophrenia spectrum disorders

We retrieved 24 human postmortem studies examining mGluR-related biology in patients with schizophrenia spectrum disorders [14–37]. Most studies did not show statistically significant group differences, while the findings varied by receptor subtype, brain region, and assay (**Figure 3**).

**Figure 3.**
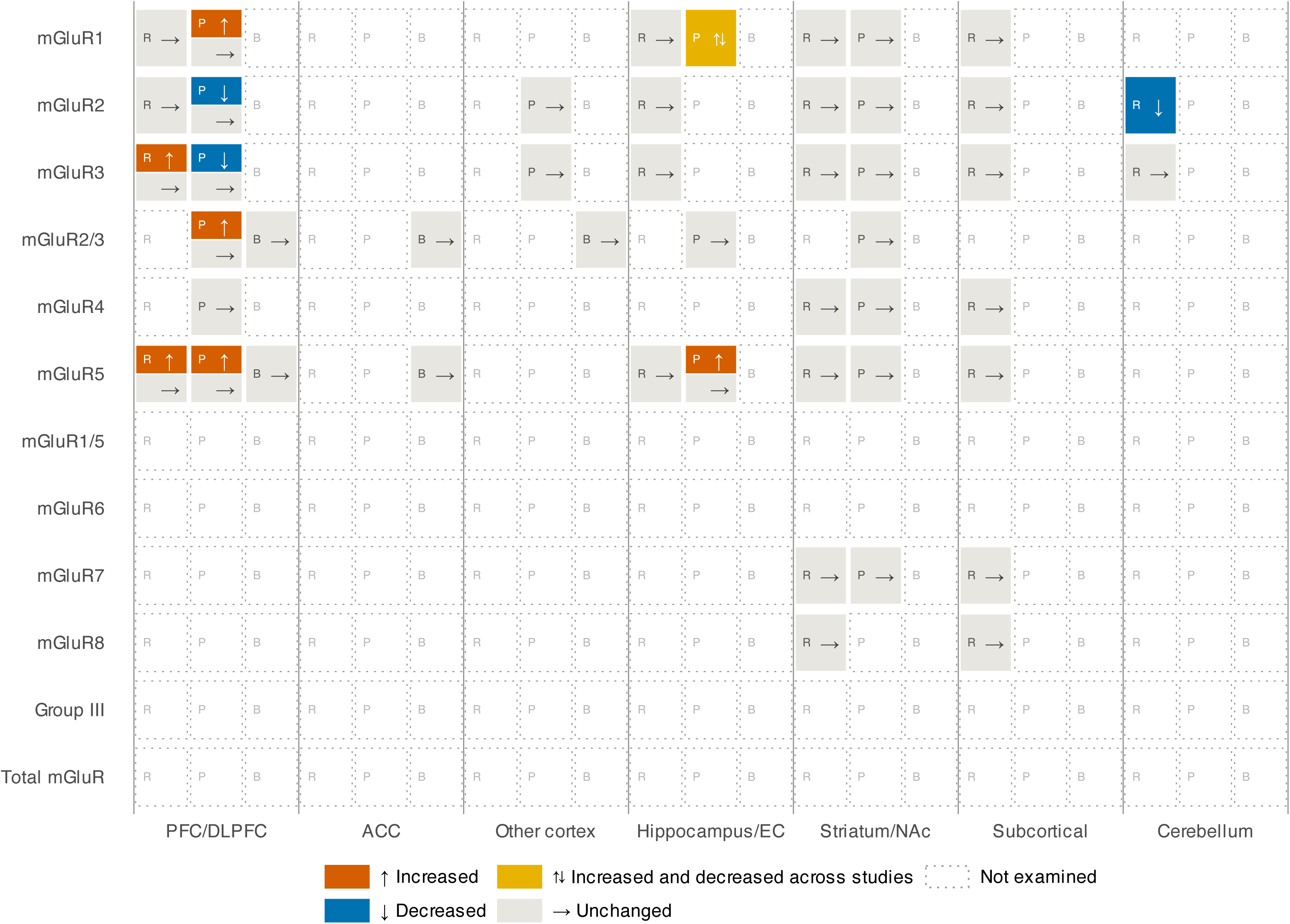
Evidence matrix of postmortem mGluR findings in schizophrenia. R, transcript abundance; P, protein abundance or immunoreactivity; B, radioligand binding. Abbreviations: ACC: Anterior Cingulate Cortex; BA: Brodmann Area; DLPFC: Dorsolateral Prefrontal Cortex; EC: Entorhinal Cortex; mGuR: Metabotropic Glutamate Receptor; NAc: Nucleus Accumbens; PFC: Prefrontal Cortex.

#### 1. Group I receptors (mGluR1 and mGluR5)

At the transcript level, *GRM5* expression was unchanged in the PFC/dorsolateral prefrontal cortex (DLPFC) [14, 15], hippocampal subfields [16, 17], and thalamic nuclei [18], and nucleus accumbens (NAc) [37]. However, Ohnuma *et al.* examined *GRM5* mRNA in PFC (BA9, BA10, and BA11) from six individuals with schizophrenia and 10 controls, showing increased *GRM5* expression in the pyramidal cell layers of BA11 [15]. With respect to protein levels, mGluR5 protein abundance was unchanged in the PFC and striatum [19], BA10 [20], DLPFC [21], and NAc [23]. Similarly, mGluR1 protein was unchanged in the striatum and NAc [19, 23].

However, mGluR1a immunoreactivity was increased across the PFC [19] and mGluR1 protein was nominally increased in the hippocampus [16]. Total mGluR5 in Cornu Ammonis 1 (CA1) was increased by 42%, accompanied by increases in its trafficking partners Norbin and Tamalin [22]. In the same CA1 tissue series, total mGluR1a was reduced by 32.7% [25]. Dimeric mGluR5 protein was increased in the DLPFC of patients with schizophrenia compared to controls [24], although unchanged DLPFC mGluR5 binding was reported in an overlapping cohort [21].

#### 2. Group II receptors (mGluR2 and mGluR3)

In studies examining mRNA, *GRM2/3* expression was unchanged in the PFC/DLPFC [15, 16, 26], hippocampus [16, 35], thalamic nuclei [18], and NAc [37]. Conversely, Bullock *et al*. compared *GRM2* and *GRM3* mRNA in lateral cerebellar hemisphere tissue from 13 males with schizophrenia and 13 matched male controls, and demonstrated that *GRM2* was reduced, whereas *GRM3* was unchanged [28]. A splice-variant analysis found modest increases in full-length and all-isoform *GRM3* measures only in the National Institute of Mental Health DLPFC cohort. Additionally, *GRM3Δ4* expression did not differ by diagnosis but was associated with the schizophrenia-risk single nucleotide polymorphism rs2228595 [29]. At the protein level, DLPFC mGluR3 was reduced in two studies [30, 31]. Dimeric/oligomeric mGluR3 was reduced in BA10 despite unchanged total mGluR3 [32]. mGluR2 was reduced regardless of antipsychotic use in one DLPFC study, in which reduced mGluR3 was confined to antipsychotic-negative cases [30], but was unchanged in the DLPFC, temporal cortex, and motor cortex in another [31].

Conversely, mGluR2/3 immunoreactivity was increased across the PFC, but not in the striatum (caudate, putamen, and NAc) [19]. At the receptor-binding level, mGluR2/3 binding was unchanged in the DLPFC [34, 36] and anterior cingulate cortex (ACC) (BA24) [27]. Using in situ hybridization, Ghose *et al*. compared mGluR3 and glutamate carboxypeptidase II transcripts along the anterior-posterior hippocampal axis in 20 individuals with schizophrenia and 20 controls [35]. mGluR3 expression did not differ in the dentate gyrus, CA1, or CA3 of either the anterior or posterior hippocampus; however, glutamate carboxypeptidase II was reduced in anterior CA1, and its normal correlation with mGluR3 in anterior CA3 was not observed in patients with schizophrenia.

#### 3. Group III receptors (mGluR4, mGluR6, mGluR7, and mGluR8)

No statistically significant alterations were reported for group III receptor expression, although coverage was limited to *GRM* transcript in thalamus [18] and NAc [37], and prefrontal and striatal mGluR4a protein [19]. No included schizophrenia study specifically assessed mGluR6.

### Mood Disorders

Twelve studies examined mGluR-related measures in patients with MDD or bipolar disorder [7, 26, 27, 29, 36, 38–44]. One additional study investigated individuals who died by suicide without a recorded psychiatric diagnosis and is also discussed in this section [45].

#### 1. Major depressive disorder

Eleven postmortem studies investigated molecular alterations in the mGluR system in patients with MDD, although most analyses yielded null results (**Figure 4**) [7, 26, 27, 36, 38–44].

**Figure 4.**
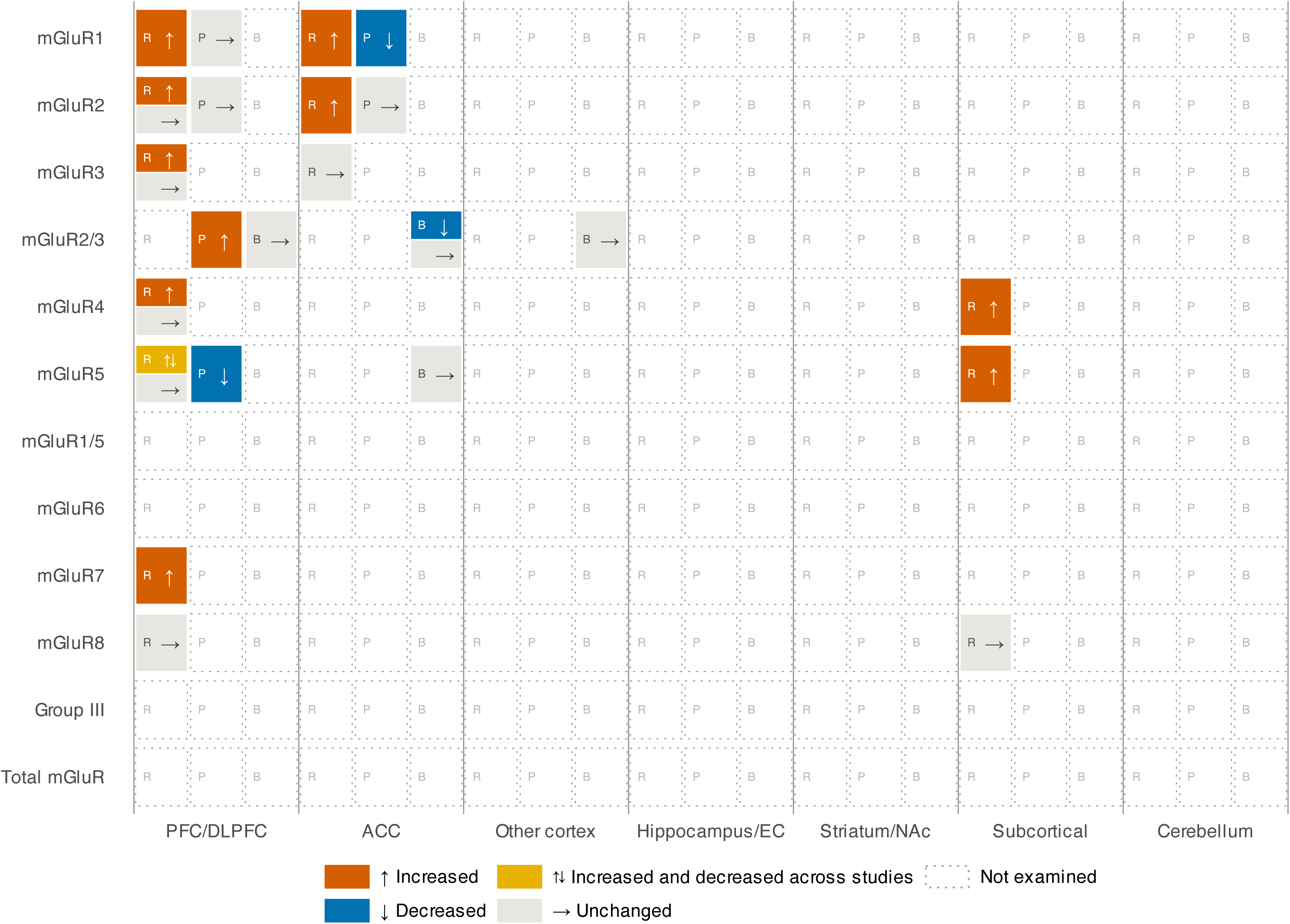
Evidence matrix of postmortem mGluR findings in major depressive disorder. R, transcript abundance; P, protein abundance or immunoreactivity; B, radioligand binding. Abbreviations: ACC: Anterior Cingulate Cortex; BA: Brodmann Area; DLPFC: Dorsolateral Prefrontal Cortex; EC: Entorhinal Cortex; mGuR: Metabotropic Glutamate Receptor; NAc: Nucleus Accumbens; PFC: Prefrontal Cortex.

At the transcript level, *GRM2/3* expression was unchanged in BA9 [26] and the DLPFC [38]. Gray *et al.* compared *GRM1, GRM2, GRM3, GRM4, GRM5,* and *GRM7* expression in BA9/46 between 53 individuals with MDD and 32 controls [43]. Overall, *GRM4* and *GRM7* were increased in the MDD group. Sex-stratified analyses showed increased *GRM1, GRM4, GRM5,* and *GRM7* in women with MDD and reduced *GRM5* in men with MDD. In locus coeruleus neurons, *GRM4* and *GRM5* expression levels were 16% and 45% higher than control values, respectively, while *GRM8* was unchanged [41]. In the BA10 pyramidal neurons, *GRM4*, *GRM5*, and *GRM8* were unchanged [41].

At the protein level, mGluR2 was unchanged in BA24 and BA46 [39]. Feyissa *et al.* compared mGluR2/3 protein in the right PFC (BA10) from 11 individuals with MDD and 11 controls and found a 67% increase in patients with MDD compared with controls [40]. In a subsequent analysis of the same cohort, mGluR5 was reduced by 32%, whereas mGluR1 was unchanged [7]. In the subgenual ACC (sgACC), mGluR1 was reduced irrespective of episode recurrence or remission status, with the reduction confirmed in an independent matched subset [42]. At the receptor-binding level, mGluR2/3 and mGluR5 binding were unchanged in BA24 [27], and DLPFC mGluR2/3 receptor density and affinity were also unchanged [38]. McOmish *et al*. noted that mGluR2/3 binding was reduced in patients with MDD in BA24 but not in BA46 or BA17 [36].

#### 2. Bipolar disorder

Five studies investigated mGluR protein, RNA, or binding in patients with bipolar disorder in addition to patients with MDD and reported predominantly null results (**Supplemental Figure S2A**) [26, 27, 29, 36, 39]. The only exception was reduced *GRM2* expression in BA9, with unchanged *GRM3* [26]. At the protein level, mGluR2 abundance was unchanged in BA24 and BA46 [39]. At the receptor-binding level, mGluR2/3 and mGluR5 binding were unchanged in BA24 [27], and mGluR2/3 binding was unchanged across BA24, BA46, and BA17 [36].

#### 3. Suicide-related findings

Suicide-related mGluR alterations were examined in five studies (**Supplemental Figure S2B**) [37, 39, 43–45]. In DLPFC (BA9/46), *GRM2* expression was higher in individuals with MDD who died by suicide (MDD-S), than in those who died from other causes (MDD-NS), whereas *GRM1*, *GRM3*, *GRM4*, *GRM5*, and *GRM7* did not differ by suicide status [43]. Conversely, another study found lower DLPFC *GRM2* expression in MDD-S than in MDD-NS, although both groups showed higher *GRM1*, *GRM2*, and *GRM3* expressions than controls [44]. In the ACC, *GRM1* and *GRM2* levels were higher in MDD-S than in both comparison groups, whereas *GRM3* was unchanged [44]. Among men who died by suicide without a documented psychiatric diagnosis or psychotropic treatment, *GRM5* level was increased by approximately 29% in the amygdala and 30% in the hippocampus but was unchanged in the DLPFC, compared with matched controls [45]. At the protein level, increased mGluR2 protein in BA46 was reported among individuals who died by suicide, regardless of diagnosis [39]. In schizophrenia, *GRM3* mRNA in the NAc was 27.7% higher in individuals with schizophrenia who died by suicide than in those who died from other causes [37].

### Neurodegenerative Diseases

#### 1. Alzheimer’s disease

Nine studies have examined mGluR in Alzheimer’s disease (AD) [46–54], and findings for mGluR5 varied by region, cell type, and molecular level (**Supplemental Figure S3A**). mGluR5 binding was 5.2-fold higher in the frontal cortex and 2.5-fold higher in the hippocampus in severe AD, but was unchanged in the cerebellum [46]. In the caudate, the proportion of mGluR5-immunopositive striatal neurons increased from approximately 40% in young controls to 80% in older controls and 92% in individuals with AD, although the difference between older controls and individuals with AD was not significant [47]. Similarly, another study found no diagnostic differences in hippocampal mGluR5 or entorhinal *GRM5* levels, although among controls hippocampal mGluR5 was positively associated with Braak stage and negatively associated with amyloid burden [48]. Cell-type-resolved analysis further showed that astrocytic mGluR5 was increased in CA1 from early Braak stages and correlated positively with both Braak stage and Aβ plaque burden, with smaller increases in CA2, CA3, the dentate gyrus, and entorhinal cortex [51].

At the transcriptomic level, female-stratified bulk RNA-sequencing analyses showed that *GRM5* expression was negatively associated with tau burden in the DLPFC but positively associated with tau burden in the posterior cingulate cortex [50]. Bartas *et al*. reported that single-nucleus analyses predicted increased glutamatergic communication involving *GRM1*, *GRM5*, and *GRM8* pathways, although *GRM1* expression itself was not increased [49]. These results represent computationally inferred communication rather than direct receptor measurements.

Regarding mGluR2, hippocampal protein abundance was increased in AD, particularly in CA1 and CA3 pyramidal neurons, and overlapped neurofibrillary pathology [52]. In neurons of the nucleus basalis of Meynert, *GRM2* level was reduced by 55% in pS422-positive pretangle-bearing neurons compared with unlabelled neurons, whereas *GRM1* and *GRM3–8* were unchanged [53]. In precuneus pyramidal neurons from the same cohort, *GRM1*, *GRM2*, and *GRM5* levels were nominally higher in AD but did not remain significant after false discovery rate (FDR) correction [54].

#### 2. Diffuse Lewy body disease

Evidence in diffuse Lewy body disease (DLBD) is limited to one postmortem study (**Supplemental Figure S3B**), in which Group I and group II mGluR binding was reduced in patients with pure-form or common-form DLBD compared with controls, whereas mGluR1 protein increased in the pure form and decreased in patients with common-form DLBD [55].

### Other Disorders

#### 1. Fragile X syndrome

Two studies have examined the density or expression of mGluRs in patients with Fragile X syndrome (FXS) and related conditions (**Supplemental Figure S4A**) [56, 57]. In the PFC of individuals with FXS or carriers, a marginal increase in mGluR5 binding density (+16%) and an increase in mGluR5 protein (+32%) were reported, compared with controls, although neither difference remained significant when the four carriers were excluded from the analysis [56]. In contrast, *GRM5* transcript and mGluR5 protein were reduced in the cerebellum of individuals with fragile X-associated tremor/ataxia syndrome or premutation carriers, compared with age-matched healthy controls, particularly with higher CGG repeat length, while mGluR1 was unchanged [57].

#### 2. Post-traumatic stress disorder

To date, only one postmortem study of mGluR has been conducted (**Supplemental Figure S4B**) [8]. Holmes *et al*. reported unchanged *GRM5* expression, alongside a 3.8-fold upregulation of *SHANK1* expression and a 3.5-fold reduction in *FKBP5* expression in the PTSD group compared with the control group [8].

#### 3. Rett syndrome

Two studies from a single sample of nine girls with Rett syndrome (RS) and ten age-matched female controls measured mGluR density without specifying subtype (**Supplemental Figure S4C**) [58, 59]. They reported no differences in mGluR density between patients with RS and controls in either the superior frontal gyrus or the basal ganglia, with only trends toward higher density in younger patients compared with controls, in the caudate nucleus (+44%), and in the globus pallidus (+31%) [58, 59]. When participants were divided into two age groups (under and over 10 years), both groups showed a significant age-related decline in receptor density.

#### 4. Autism spectrum disorder

Five studies have investigated mGluR in individuals with autism spectrum disorder (ASD) (**Supplemental Figure S5**) [60–64]. Dimerized and total mGluR5 were increased in the superior frontal gyrus and in the cerebellar vermis in children with ASD compared with controls, with no difference in adults [60, 64]. In adults, mGluR3 protein was reduced in BA9 [61] and *GRM5* expression was reduced in the DLPFC in a separate cohort [63].

#### 5. Alcohol use disorder

Four studies examined mGluR-related measures in alcohol use disorder (AUD) (**Supplemental Figure S6**) [65–68], three of which used the same donor series of 17 individuals with AUD, comprising Cloninger type 1 and type 2 subgroups, and ten controls [65–67]. In the CA2, mGluR1/5 binding was increased in patients with type 1 AUD compared with patients with type 2 AUD and controls [65]. Additionally, mGluR1/5 binding was also increased in the anterior insula, putamen, and head of the caudate nucleus in both patients with type 1 and with type 2 AUD, compared with controls, and in the hippocampus in type 2 AUD [66].

mGluR2/3 binding was higher in the pregenual ACC in individuals with type 2 AUD compared to controls [67]. Using RNA sequencing of postmortem hippocampal tissue, Enoch *et al.* found increased *GRM4* expression in patients with AUD compared with the combined control and cocaine-use-disorder groups after FDR correction; however, this pooled comparison limits direct comparison with findings from other AUD studies [68].

### Medication exposure

Nineteen of the 57 included studies examined associations between medication exposure and an mGluR outcome measure in human tissue, indexing exposure either as cumulative or lifetime dose, or as drug status and estimated receptor occupancy at the time of death (**Table 1**).

Overall, antipsychotic exposure was not associated with most measures of mGluR protein and *GRM* transcript levels or mGluR binding in schizophrenia. Postmortem studies have reported no association with cumulative antipsychotic dose for mGluR3 immunoreactivity in the superior temporal cortex [33], mGluR2/3 binding in the DLPFC (BA46) [34], mGluR5 binding and protein in the DLPFC [21, 24], mGluR2/3 and mGluR5 binding in the ACC [27], mGluR1 protein in CA1 [25], mGluR1 and mGluR5 protein in the NAc [23], or *GRM2/3* in the frontal cortex (BA9) [26]. Similarly, the *GRM3* increase in the DLPFC was independent of daily and lifetime doses [29]. Qualitatively, mGluR1a, mGluR2/3, mGluR4 and mGluR5 protein in the PFC and striatum did not differ between donors who had received antipsychotics and donors who had not [19]. However, several region– and receptor-specific associations were reported. In CA1, mGluR5 protein correlated negatively with lifetime chlorpromazine equivalents [22]. In the PFC (BA10), but not the DLPFC (BA46), *GRM5* expression correlated with lifetime antipsychotic dose, in the absence of a diagnostic difference in either region [14]. Dela Cuesta-Barrutia *et al*. reported that the reduction in mGluR3 protein and the 64% increase in *GRM3* expression were both confined to donors who were not on antipsychotics, whereas the reduction in mGluR2 protein was present irrespective of medication exposure. Estimated 5-HT2A receptor occupancy correlated positively with mGluR3 protein [30]. In the NAc, *GRM4* expression was significantly increased in patients with schizophrenia with a history of antidepressant use [37].

In mood disorders, GRM2/3 expression and mGluR2/3 binding in the PFC were unchanged in donors on antidepressants [38]. *GRM1-5* and *GRM7* transcript levels in the DLPFC (BA9/46) [43], *GRM2/3* in the frontal cortex (BA9) [26], and mGluR1 protein in the subgenual ACC [42] were not associated with antidepressant status. For antipsychotics, the reduction in mGluR1 protein also persisted after adjustment for exposure [42].

In other disorders, *GRM5* and *GRM8* levels in the ACC (BA24) and PFC (BA10) did not differ between drug-exposed and unexposed donors with ASD [62], nor did *GRM5* mRNA in the subgenual ACC (BA25) differ between donors who were and were not medicated at the time of death in PTSD [8].

## DISCUSSION

Although findings across studies were heterogeneous and often inconsistent, mGluR-related alterations may be observed in some neuropsychiatric disorders and neurodegenerative disorders. Among the mGluR subtypes, mGluR5 and mGluR2/3 were extensively investigated, followed by mGluR1, whereas evidence for the remaining group III receptors was comparatively limited. Regarding brain regions, studies mainly investigated the PFC, ACC, and hippocampus, whereas other cortical and subcortical regions were investigated less frequently.

### Cross-disorder receptor patterns

mGluR5 and mGluR1 constitute group I mGluRs and are Gq/11-coupled receptors that activate phospholipase Cβ, leading to downstream Ca²⁺ mobilization and protein kinase C signaling [2]. mGluR5 is widely expressed in the cortex, hippocampus, and striatum, and predominantly localized at postsynaptic and perisynaptic sites [69]. Through interactions with Homer and Shank scaffolding proteins, mGluR5 is incorporated into postsynaptic signaling complexes linked to inositol trisphosphate receptors and NMDA receptor-associated proteins and contributes to glutamatergic signaling and synaptic plasticity [2, 70]. Among the available postmortem mGluR studies, mGluR5 was investigated most extensively; however, no consistent direction of difference was evident across disorders or measurement modalities. In MDD, mGluR5 protein was reduced in BA10 but *GRM5* expression was increased in locus coeruleus neurons and showed sex-dependent changes in the DLPFC [7, 41, 43]. In AD, mGluR5 alterations may be linked to neuropathological burden in a region– and cell type-dependent manner, with *GRM5* expression associated with tau pathology in cortical regions and astrocytic mGluR5 protein level associated with Aβ burden in the hippocampus [50, 51]. Other findings included increased GRM5 mRNA in the amygdala and hippocampus of individuals who died by suicide [45]; increased mGluR5 protein in the cerebellar vermis and superior frontal cortex (BA9) of children with ASD [60, 64]; increased mGluR5 protein in the PFC of individuals with FXS or carriers, but decreased mGluR5 protein in the cerebellum of patients with fragile X-associated tremor/ataxia syndrome [56, 57]; and increased mGluR1/5 binding in CA2, putamen, caudate head, and anterior insula of patients with AUD [65, 66]. In schizophrenia, increased mGluR5 protein and its trafficking partners Norbin and Tamalin were observed in CA1 [22], despite predominantly null findings in the PFC, DLPFC, ACC, NAc, and other hippocampal analyses [14, 17, 19, 21, 23, 27]. The accompanying alterations in Norbin, Tamalin, receptor dimerization, and *SHANK1* suggest that mGluR-related abnormalities may reflect a broader, multilevel disturbance of the glutamatergic system, arising from the interplay among receptor abundance, trafficking, anchoring, dimerization, and synaptic organization [8, 22, 24, 32, 61].

Group II mGluRs (mGluR2 and mGluR3) are Gi/o-coupled receptors that inhibit adenylyl cyclase and modulate glutamatergic transmission [2]. mGluR2 is expressed mainly at presynaptic neuronal terminals, and suppresses glutamate transmission, whereas mGluR3 is more broadly distributed across presynaptic and postsynaptic neuronal compartments and glial cells, particularly astrocytes, where it can engage neuroprotective signaling, including the production and release of transforming growth factor-β [3]. According to available postmortem studies, in patients with schizophrenia, reduced mGluR2 or mGluR3 protein was reported most consistently in the DLPFC [30, 31], whereas transcript and binding studies in the DLPFC, ACC, and other regions have yielded largely null or inconsistent findings [18, 26, 27, 29, 33, 34]. In individuals with MDD, mGluR2/3 protein was increased in BA10 [40], while binding was reduced in BA24 and unchanged in other frontal regions [27, 36, 38]. In AD, mGluR2 protein was increased in vulnerable hippocampal pyramidal neurons, whereas *GRM2* was reduced specifically in pretangle-bearing nucleus basalis neurons [52, 53].

Overall, mGluR alterations show no uniform direction across neuropsychiatric disorders and may vary with diagnostic status, disease severity, pathological context, and brain region.

### Brain-region specificity

The PFC and ACC were the most frequently examined cortical regions. Most studies, however, examined individual Brodmann areas or subregions separately using different receptor measures and methodologies, limiting direct assessment of anatomical variation within these regions. In schizophrenia, increased neuronal *GRM5* mRNA was reported in BA11 [15], whereas findings in BA9, BA10, and BA46 were largely null [14, 19–21]. In MDD, BA10 showed increased mGluR2/3 and reduced mGluR5 protein [7, 40], while reduced mGluR1 was reported in the sgACC and reduced mGluR2/3 binding in BA24 [36, 42].

Hippocampal findings similarly varied according to receptor subtype, subfield, and disease context. In schizophrenia, CA1 showed increased mGluR5 together with Norbin and Tamalin, but reduced mGluR1 [22, 25], whereas findings in other hippocampal subfields and transcript-level analyses were largely null [16, 17]. In AD, increased mGluR2 was reported in vulnerable CA1 and CA3 pyramidal neurons and overlapped with neurofibrillary pathology [52]. Findings for mGluR5 were more heterogeneous across molecular measures and pathological contexts, with increased hippocampal binding reported in some studies [46], no diagnostic difference in others [48], and more recent evidence of increased astrocytic mGluR5 in CA1 that correlated with Braak stage and Aβ plaque burden [51]. AUD studies identified subtype-specific alterations in hippocampal mGluR1/5 binding [65, 66].

Taken together, mGluR abnormalities may vary across cortical areas, hippocampal subfields, or cell types. However, uneven regional sampling and few comparable multi-region analyses leave it unclear whether findings from individual Brodmann areas or hippocampal subfields can be generalized to the broader regions.

### Cross-modality concordance and discordance

Some studies applied multiple assays to the same specimens, whereas others assessed different measurement modalities and/or brain regions in separate publications from the same donor series. Within-study, shared-cohort, and independent-cohort comparisons revealed concordant and discordant patterns.

Concordance comprised both parallel molecular alterations and agreement in null findings (**Table 2B**) [21, 38, 48, 63]. However, discordance was more frequent among within-study cross-level comparisons (**Table 2A**), including receptor protein alterations despite unchanged transcript abundance in several schizophrenia studies and disagreement between protein abundance and radioligand binding in DLBD [16, 21, 24, 30, 55]. Notably, increased *GRM3* expression coexisted with reduced mGluR3 immunoreactivity in antipsychotic-free patients [30]. Similarly, across two reports based on the same BA24 donor series, reduced [^3^H]LY341495 binding coexisted with unchanged GRM2 protein; the authors argued that the reduction in binding was therefore attributable to lower mGluR3 rather than mGluR2 [36, 39]. Across independent AD hippocampal cohorts, increased mGluR5 binding [46] and increased mGluR5 immunoreactivity in CA1 astrocytes [51] were reported, whereas bulk immunofluorescence in a larger cohort showed no overall difference [48].

Therefore, transcript abundance, receptor protein, and radioligand binding capture distinct but interconnected levels of receptor biology. Transcript measurements reflect steady-state RNA abundance, whereas protein abundance is additionally shaped by translation, degradation, assembly, trafficking, and cellular or subcellular localization. Accordingly, RNA–protein correspondence varies across genes and brain regions [71, 72]. Radioligand binding, in contrast, reflects ligand-accessible receptor sites and is influenced by receptor affinity, conformation, localization, and ligand subtype selectivity [73]. For example, the opposing *GRM3* transcript and mGluR3 immunoreactivity findings may be compatible with compensatory transcriptional responses or altered translation or protein turnover, although the available data cannot distinguish among these mechanisms [30]. Similarly, discordance between protein abundance and radioligand binding may indicate differential changes in total and ligand-accessible receptor pools or, for ligands lacking complete subtype selectivity, the masking of subtype-specific alterations. The AD findings further suggest that alterations confined to particular cell types or receptor pools may be diluted in bulk-tissue measurements [51]. Although cross-modality discordance does not establish the underlying mechanism, integrating transcript, protein, cell-specific, and ligand-binding measurements, ideally in the same specimens or across matched postmortem cohorts and complementary PET studies, may provide a more comprehensive understanding of mGluR dysregulation as a multilevel system.

### Potential effects of medication exposure on mGluR alterations

Medication exposure was not uniformly related to mGluR measures. Rather, the available evidence suggests that its influence may vary across receptor subtype, brain region, and treatment context.

Lifetime antipsychotic exposure correlated negatively with mGluR5 protein in CA1 despite increased mGluR5 protein in patients with schizophrenia [22]. Similarly, mGluR3 protein in the DLPFC was reduced in antipsychotic-negative donors but not in antipsychotic-positive donors and estimated 5-HT2A receptor occupancy correlated positively with mGluR3 protein level [30]. Antipsychotic exposure may therefore attenuate or mask disease-associated abnormalities, although the two studies indexed exposure over different time windows, with one reflecting cumulative lifetime dose and the other exposure at the time of death. In addition, lifetime antipsychotic dose was positively associated with *GRM5* in PFC (BA10) despite no case-control difference [14], suggesting an antipsychotic medication-related alteration; however, it is unclear whether this association reflects normalization of a potential disease-related abnormality.

Accordingly, medication exposure should neither be ignored nor treated as a uniform explanation for case-control differences. Treatment could attenuate or mask some mGluR abnormalities, but the evidence is insufficient to establish the net direction or magnitude of its effect. Since these findings are observational, cumulative exposure may vary with illness severity, duration, treatment response, and comorbidity, creating confounding by indication.

### Therapeutic implications: what postmortem data can and cannot support

In clinical research, mGluRs have been investigated as therapeutic targets for orthosteric agonists, antagonists, and allosteric modulators [5]. Although several mGluR modulators have entered clinical development, none has established a role in standard treatment of neuropsychiatric disorders. In schizophrenia, the mGluR2/3 agonist prodrug pomaglumetad methionil did not show significant overall efficacy in a meta-analysis of randomized trials [74]. In MDD, trials of the mGluR5 modulator basimglurant and the group II modulators decoglurant and MK-1942 did not establish antidepressant efficacy [75–77]. In AUD, the mGluR5 negative allosteric modulator GET 73 failed to reduce cue-induced craving or laboratory alcohol self-administration in hospitalized patients with AUD [78].

Although postmortem data are informative for therapeutic target nomination and anatomical localization [79], they cannot determine whether an observed alteration is causal or compensatory, which direction of receptor modulation is required, or how the target behaves within the intact and dynamic biological system. Therefore, integrating postmortem evidence with in vivo target-engagement measures and prospective biomarker-based patient stratification is warranted.

### Relationship to PET studies

Whereas postmortem studies provide a single terminal measurement and depend on donor availability, PET enables repeated within-person measurements. In vivo PET studies may therefore complement postmortem findings by examining the living brain and preserving its anatomical and systems-level context. PET measurements may also be sensitive to physiological state, and this sensitivity may provide biologically relevant information but may also introduce confounding. For instance, food intake and plasma glucose were associated with changes in [¹¹C]ABP688 binding in healthy individuals [80]. Longitudinal designs and pharmacological interventions can help examine within-person changes in receptor availability in relation to symptoms, disease stage, and treatment response [81, 82]. Human mGluR PET research has predominantly focused on mGluR5. In schizophrenia, two studies using [¹¹C]ABP688 found no differences in mGluR5 availability compared to healthy controls, although lower availability in the temporal cortex and caudate was associated with more severe negative symptoms and poorer cognitive or social functioning [83, 84]. In MDD, several studies of younger or unmedicated cohorts reported lower mGluR5 availability [7, 85, 86], whereas no difference was shown in studies of late-life MDD and another larger [^18^F]FPEB cohort [87, 88]. Recently, a longitudinal study reported lower baseline mGluR5 availability in patients with MDD and increases in the DLPFC and ventromedial PFC after eight weeks of vortioxetine treatment, which correlated with symptomatic improvement [89].

PET findings show some convergence with the present postmortem findings, while the two approaches capture different aspects of receptor biology. In schizophrenia, the unchanged mGluR5 availability compared with controls is broadly consistent with several postmortem studies reporting no alterations in mGluR5 binding or total protein [19–21], although alterations in receptor dimerization or regulatory proteins might not be fully captured in PET ligand availability [24]. Since mGluR2/3 has been little examined with PET in patients with schizophrenia, future mGluR2/3 imaging could help assess the in vivo relevance of these postmortem findings. In MDD, Deschwanden *et al*. reported concordant reductions in mGluR5 PET availability and mGluR5 protein in BA10 [7], whereas subsequent null findings and longitudinal changes suggest that mGluR5 availability may also depend on age, clinical state, and treatment [87–89]. In AD, studies using [^11^C]ABP688, [^18^F]FPEB, and [^18^F]PSS232 have shown largely consistent reductions centered on the hippocampus and adjacent medial temporal regions [90–92]. This pattern contrasts with heterogeneous postmortem findings and may reflect disease stage, atrophy or synaptic loss, cell-type composition, or differences between ligand-accessible and total receptor measures [46, 48, 51]. Therefore, PET may provide a useful translational bridge for assessing whether and when targets nominated by postmortem studies remain available in vivo.

### Limitations

This review has several limitations. First, many disorders, receptor subtypes, and brain regions were represented by only a small number of studies, and several reports used the same or overlapping donor cohorts. Thus, publication counts may overstate independent evidence. Second, postmortem findings cannot be assumed to represent receptor status in the living brain. Molecular measures may be affected by agonal conditions, postmortem interval, tissue pH, RNA integrity, tissue handling, and other pre– and postmortem factors [93]. Cross-sectional postmortem comparisons also cannot establish whether an observed alteration preceded illness or instead reflected illness duration, treatment or substance exposure, comorbidity, terminal events, or neurodegeneration. In addition, the limited availability of postmortem brain tissue constrains sample sizes and generally precludes repeated within-person assessment over time. Third, clinical information, including symptom severity, illness stage and duration, mood state, lifetime medication exposure, smoking, and other substance use, was incompletely and inconsistently reported across the primary studies. We therefore could not reliably relate mGluR abnormalities to clinical phenotypes or distinguish trait-related changes from state-, treatment-, or exposure-related effects. Fourth, a quantitative meta-analysis was not feasible because studies differed substantially in diagnosis, brain region, receptor subtype, tissue fraction, assay, and outcome reporting, and many did not provide comparable effect estimates or variance measures. Moreover, mRNA abundance, total receptor protein, immunoreactivity, receptor dimerization, and radioligand binding represent distinct biological or measurement levels and should not be treated as equivalent indicators of receptor abundance or function [94, 95]. Accordingly, the findings should be regarded as provisional, region– and assay-specific signals rather than evidence of a unified cross-disorder mGluR abnormality.

### Future directions

It would be useful for future research to build on findings supported by independent postmortem cohorts and, where possible, convergent evidence across molecular assays. PET tracers targeting mGluR provide an opportunity for cross-modal triangulation of postmortem findings with in vivo receptor availability [96].

In schizophrenia-spectrum disorders, DLPFC mGluR2/3 alterations may merit further investigation [30]. PET studies could define the DLPFC as a primary region of interest and compare antipsychotic-naïve individuals with first-episode psychosis and chronically treated patients to explore potential effects of illness stage and treatment. In MDD, mGluR5 has been examined with both postmortem and PET approaches, and longitudinal treatment-related changes were also evaluated [7, 89]. Longitudinal designs with repeated imaging during treatment could further characterize changes in mGluR availability and their relationship with clinical response.

In ASD, developmental stage may be an important consideration. Increased mGluR5 measures in childhood tissue and reduced *GRM3* or *GRM5* expression in adult cortical samples raise the possibility that alterations vary across development. Studies with larger samples spanning developmental stages may help clarify these patterns. In AD and MCI, an important question is how mGluR abnormalities relate to amyloid and tau pathology, synaptic loss, and neurodegeneration. Recent region– and cell-specific findings suggest mGluR5 alterations may depend on neuropathological burden and cellular context. Multimodal longitudinal studies combining mGluR5 imaging with amyloid, tau, synaptic density, and structural neurodegeneration may help clarify the temporal relationships among these processes [91]. In AUD, independent replication may be particularly informative because several reports may have been based on the same or overlapping donor cohorts [65–67].

More broadly, single-cell and spatial transcriptomics may help resolve limitations of bulk-tissue mRNA studies by distinguishing neuronal, glial, and layer-specific receptor changes. Spatial proteomics and receptor-specific autoradiography may further complement conventional protein measurements, which do not fully capture receptor localization or molecular organization such as dimerization. Where feasible, integration of postmortem findings with PET, including harmonized imaging–brain-bank cohorts or PET-to-autopsy studies, may help relate molecular alterations identified ex vivo to receptor availability in vivo. Together with transparent reporting of overlapping donor cohorts, these approaches may facilitate interpretation of both convergent and divergent findings across studies. A combination of these strategies may help clarify the biological and clinical significance of mGluR alterations and the factors underlying heterogeneity across studies, which will provide a more refined basis for future translational investigation.

## CONCLUSION

Postmortem studies suggested mGluR-related alterations in some neuropsychiatric and neurodegenerative disorders, although findings varied substantially across receptor subtypes, brain regions, molecular measures, and clinical populations. Overall, these findings may reflect context-dependent molecular signals rather than evidence of uniform or disease-specific mGluR abnormalities. Further integration with in vivo imaging and other complementary approaches may help clarify the biological and clinical significance of both convergent and divergent findings.

### Funding

This work received no specific funding.

Supplementary information is available at MP’s website.

### Financial support

This work received no specific funding.

### Previous presentation

None.

## Acknowledgments

None.

## Use of AI-assisted technologies in the writing process

In the writing of this manuscript, the authors used AI for proofreading the manuscript. The authors have reviewed the content and take full responsibility for the content of the publication.

## ABBREVIATIONS

Aβ: Amyloid-beta
ACC: Anterior Cingulate Cortex
AD: Alzheimer’s Disease
ASD: Autism Spectrum Disorder
AUD: Alcohol Use Disorder
BA: Brodmann Area
CA: Cornu Ammonis
DLBD: Diffuse Lewy Body Disease
DLPFC: Dorsolateral Prefrontal Cortex
FDR: False Discovery Rate
FXS: Fragile X Syndrome
GRM: Glutamate Metabotropic Receptor
MCI: Mild Cognitive Impairment
MDD: Major Depressive Disorder
mGluR: Metabotropic Glutamate Receptor
NAc: Nucleus Accumbens
PET: Positron Emission Tomography
PFC: Prefrontal Cortex
PTSD: Post-Traumatic Stress Disorder
RS: Rett Syndrome

