## Supplemental materials for "Postmortem Alterations of Metabotropic Glutamate Receptors across Neuropsychiatric Disorders: A Systematic Review"

**Table of contents**

**Supplemental Table S1.	PRISMA 2020 Checklist of items to include when reporting a systematic review involving a meta-analysis**

**(Pages 2–6)**

**Supplemental Table S2.	The search strategy of MEDLINE via PubMed (Page 7)**

**Supplemental Table S3.	The search strategy of Web of Science (Page 8)**

**Supplemental Table S4.	The search strategy of EMBASE via Ovid (Page 9)**

**Supplemental Table S5.	Eligibility criteria of studies (Page 10)**

**Supplemental Figure S1.	Risk of bias of studies included in the review (Page 11-12)**

**Supplemental Figure S2.	Evidence matrix of postmortem mGluR findings in (A) Bipolar disorder and (B) Patients who died by suicide
(Pages 13-14)**

**Supplemental Figure S3.	Evidence matrix of postmortem mGluR findings in (A) Alzheimer’s disease and (B) Diffuse Lewy Body Disease. (Pages 15-16)**

**Supplemental Figure S4.	Evidence matrix of postmortem mGluR findings in (A) Fragile X-related conditions, (B) Rett syndrome, (C) Post-traumatic stress disorder (Pages 17-18)**

**Supplemental Figure S5.	Evidence matrix of postmortem mGluR findings in Autistic spectrum disorder (Pages 19-20)**

**Supplemental Figure S6.	Evidence matrix of postmortem mGluR findings in Alcohol use disorder (Pages 21-22)**

**Supplementarl Table S1.	PRISMA 2020 Checklist of items to include when reporting a systematic review.**

| **Section and Topic** | **Item #** | **Checklist item** | **Location where item is reported** |
| --- | --- | --- | --- |
| **TITLE** | | |  |
| Title | 1 | Identify the report as a systematic review. | 1 |
| **ABSTRACT** | | |  |
| Abstract | 2 | See the PRISMA 2020 for Abstracts checklist. | 2 |
| **INTRODUCTION** | | |  |
| Rationale | 3 | Describe the rationale for the review in the context of existing knowledge. | 4, 5 |
| Objectives | 4 | Provide an explicit statement of the objective(s) or question(s) the review addresses. | 5 |
| **METHODS** | | |  |
| Eligibility criteria | 5 | Specify the inclusion and exclusion criteria for the review and how studies were grouped for the syntheses. | 6 |
| Information sources | 6 | Specify all databases, registers, websites, organisations, reference lists and other sources searched or consulted to identify studies. Specify the date when each source was last searched or consulted. | 6 |
| Search strategy | 7 | Present the full search strategies for all databases, registers and websites, including any filters and limits used. | Table S2-4 |
| Selection process | 8 | Specify the methods used to decide whether a study met the inclusion criteria of the review, including how many reviewers screened each record and each report retrieved, whether they worked independently, and if applicable, details of automation tools used in the process. | 6 |
| Data collection process | 9 | Specify the methods used to collect data from reports, including how many reviewers collected data from each report, whether they worked independently, any processes for obtaining or confirming data from study investigators, and if applicable, details of automation tools used in the process. | 6 |
| Data items | 10a | List and define all outcomes for which data were sought. Specify whether all results that were compatible with each outcome domain in each study were sought (e.g. for all measures, time points, analyses), and if not, the methods used to decide which results to collect. | 6 |
|  | 10b | List and define all other variables for which data were sought (e.g. participant and intervention characteristics, funding sources). Describe any assumptions made about any missing or unclear information. | 6 |
| Study risk of bias assessment | 11 | Specify the methods used to assess risk of bias in the included studies, including details of the tool(s) used, how many reviewers assessed each study and whether they worked independently, and if applicable, details of automation tools used in the process. | 6 |
| Effect measures | 12 | Specify for each outcome the effect measure(s) (e.g. risk ratio, mean difference) used in the synthesis or presentation of results. | Not Reported |
| Synthesis methods | 13a | Describe the processes used to decide which studies were eligible for each synthesis (e.g. tabulating the study intervention characteristics and comparing against the planned groups for each synthesis (item #5)). | 6 |
|  | 13b | Describe any methods required to prepare the data for presentation or synthesis, such as handling of missing summary statistics, or data conversions. | 6 |
|  | 13c | Describe any methods used to tabulate or visually display results of individual studies and syntheses. | 6 |
|  | 13d | Describe any methods used to synthesize results and provide a rationale for the choice(s). If meta-analysis was performed, describe the model(s), method(s) to identify the presence and extent of statistical heterogeneity, and software package(s) used. | 7 |
|  | 13e | Describe any methods used to explore possible causes of heterogeneity among study results (e.g. subgroup analysis, meta-regression). | Not Reported |
|  | 13f | Describe any sensitivity analyses conducted to assess robustness of the synthesized results. | Not Reported |
| Reporting bias assessment | 14 | Describe any methods used to assess risk of bias due to missing results in a synthesis (arising from reporting biases). | Not Reported |
| Certainty assessment | 15 | Describe any methods used to assess certainty (or confidence) in the body of evidence for an outcome. | Not Reported |
| **RESULTS** | | |  |
| Study selection | 16a | Describe the results of the search and selection process, from the number of records identified in the search to the number of studies included in the review, ideally using a flow diagram. | Figure 1 |
|  | 16b | Cite studies that might appear to meet the inclusion criteria, but which were excluded, and explain why they were excluded. | Not Reported |
| Study characteristics | 17 | Cite each included study and present its characteristics. | Table 1 |
| Risk of bias in studies | 18 | Present assessments of risk of bias for each included study. | Figure S1 |
| Results of individual studies | 19 | For all outcomes, present, for each study: (a) summary statistics for each group (where appropriate) and (b) an effect estimate and its precision (e.g. confidence/credible interval), ideally using structured tables or plots. | Table 1 |
| Results of syntheses | 20a | For each synthesis, briefly summarise the characteristics and risk of bias among contributing studies. | Not Reported |
|  | 20b | Present results of all statistical syntheses conducted. If meta-analysis was done, present for each the summary estimate and its precision (e.g. confidence/credible interval) and measures of statistical heterogeneity. If comparing groups, describe the direction of the effect. | Not Reported |
|  | 20c | Present results of all investigations of possible causes of heterogeneity among study results. | Not Reported |
|  | 20d | Present results of all sensitivity analyses conducted to assess the robustness of the synthesized results. | Not Reported |
| Reporting biases | 21 | Present assessments of risk of bias due to missing results (arising from reporting biases) for each synthesis assessed. | Figure S1 |
| Certainty of evidence | 22 | Present assessments of certainty (or confidence) in the body of evidence for each outcome assessed. | Not Reported |
| **DISCUSSION** | | |  |
| Discussion | 23a | Provide a general interpretation of the results in the context of other evidence. | 17-27 |
|  | 23b | Discuss any limitations of the evidence included in the review. | 25 |
|  | 23c | Discuss any limitations of the review processes used. | 25 |
|  | 23d | Discuss implications of the results for practice, policy, and future research. | 26-27 |
| **OTHER INFORMATION** | | |  |
| Registration and protocol | 24a | Provide registration information for the review, including register name and registration number, or state that the review was not registered. | 5 |
|  | 24b | Indicate where the review protocol can be accessed, or state that a protocol was not prepared. | 5 |
|  | 24c | Describe and explain any amendments to information provided at registration or in the protocol. | 5 |
| Support | 25 | Describe sources of financial or non-financial support for the review, and the role of the funders or sponsors in the review. | 29 |
| Competing interests | 26 | Declare any competing interests of review authors. | 29 |
| Availability of data, code and other materials | 27 | Report which of the following are publicly available and where they can be found: template data collection forms; data extracted from included studies; data used for all analyses; analytic code; any other materials used in the review. | Not Reported |

**Supplemental Table S2.	The search strategy of MEDLINE via PubMed**

| #1 | "metabotropic glutamate receptor*" |
| --- | --- |
| #2 | “mGluR*” |
| #3 | “GRM*” |
| #4 | #1 OR #2 OR #3 |
| #5 | "Receptors, Metabotropic Glutamate"[Mesh] |
| #6 | #4 OR #5 |
| #7 | “post-mortem"[All Fields] |
| #8 | "postmortem"[All Fields] |
| #9 | "post mortem"[All Fields] |
| #10 | “autopsy”[All Fields] |
| #11 | “autopsied”[All Fields] |
| #12 | "brain bank*" |
| #13 | "human brain tissue" |
| #14 | #7 OR #8 OR #9 OR #10 OR #11 OR #12 OR #13 |
| #15 | "Autopsy"[Mesh] |
| #16 | "Postmortem Changes"[Mesh] |
| #17 | “Tissue Banks"[Mesh] |
| #18 | #15 OR #16 OR #17 |
| #19 | #14 OR #18 |
| #20 | "Humans"[Mesh] |
| #21 | #4 AND #19 AND #20 |

**Supplemental Table S3.	The search strategy of Web of Science**

| #1 | TS=(metabotropic glutamate receptor*) |
| --- | --- |
| #2 | TS=(mGluR*) |
| #3 | TS=(GRM*) |
| #4 | #1 OR #2 OR #3 |
| #5 | TS=(post-mortem) |
| #6 | TS=(postmortem) |
| #7 | TS=(post mortem) |
| #8 | TS=(autopsy) |
| #9 | TS=(autopsied) |
| #10 | TS=(brain bank) |
| #11 | TS=(human brain tissue) |
| #12 | #5 OR #6 OR #7 OR #8 OR #9 OR #10 OR #11 |
| #13 | TS=(human OR humans) |
| #14 | #4 AND #12 AND #13 |

**Supplemental Table S4.	The search strategy of EMBASE via Ovid**

| #1 | 'metabotropic glutamate receptor'/exp |
| --- | --- |
| #2 | 'metabotropic glutamate receptor*':ti,ab |
| #3 | mglur*:ti,ab |
| #4 | grm*:ti,ab |
| #5 | 1 or 2 or 3 or 4 |
| #6 | 'postmortem examination'/exp |
| #7 | 'postmortem':ti,ab |
| #8 | 'post mortem':ti,ab |
| #9 | autopsy:ti,ab |
| #10 | autopsied:ti,ab |
| #11 | 'brain bank*':ti,ab |
| #12 | 'human brain tissue':ti,ab |
| #13 | 'tissue bank'/exp |
| #14 | 'postmortem change'/exp |
| #15 | 6 or 7 or 8 or 9 or 10 or 11 or 12 or 13 or 14 |
| #16 | [humans]/lim |
| #17 | 5 and 15 and 16 |

**Supplemental Table S5.	Eligibility criteria of studies**

| Inclusion criteria | |
| --- | --- |
| 1. | Studies utilizing human postmortem brain tissues |
| 2. | Studies focusing on individuals diagnosed with neuropsychiatric disorders |
| 3. | Studies investigating mGluR expression, receptor binding, or mGluR-related transcripts |
| Exclusion criteria | |
| 1. | Studies using non-human animal models or tissues |
| 2. | Studies measuring the concentrations of substances acting on mGluRs (e.g., agonists, antagonists, or allosteric modulators) |
| 3. | Studies targeting downstream signaling molecules of the mGluR cascade |
| 4. | Studies being published as reviews, methodological studies, or conference abstracts |

**Supplemental Figure S1.	Risk of bias of studies included in the review.**

(A) Distribution of Overall Risk of Bias

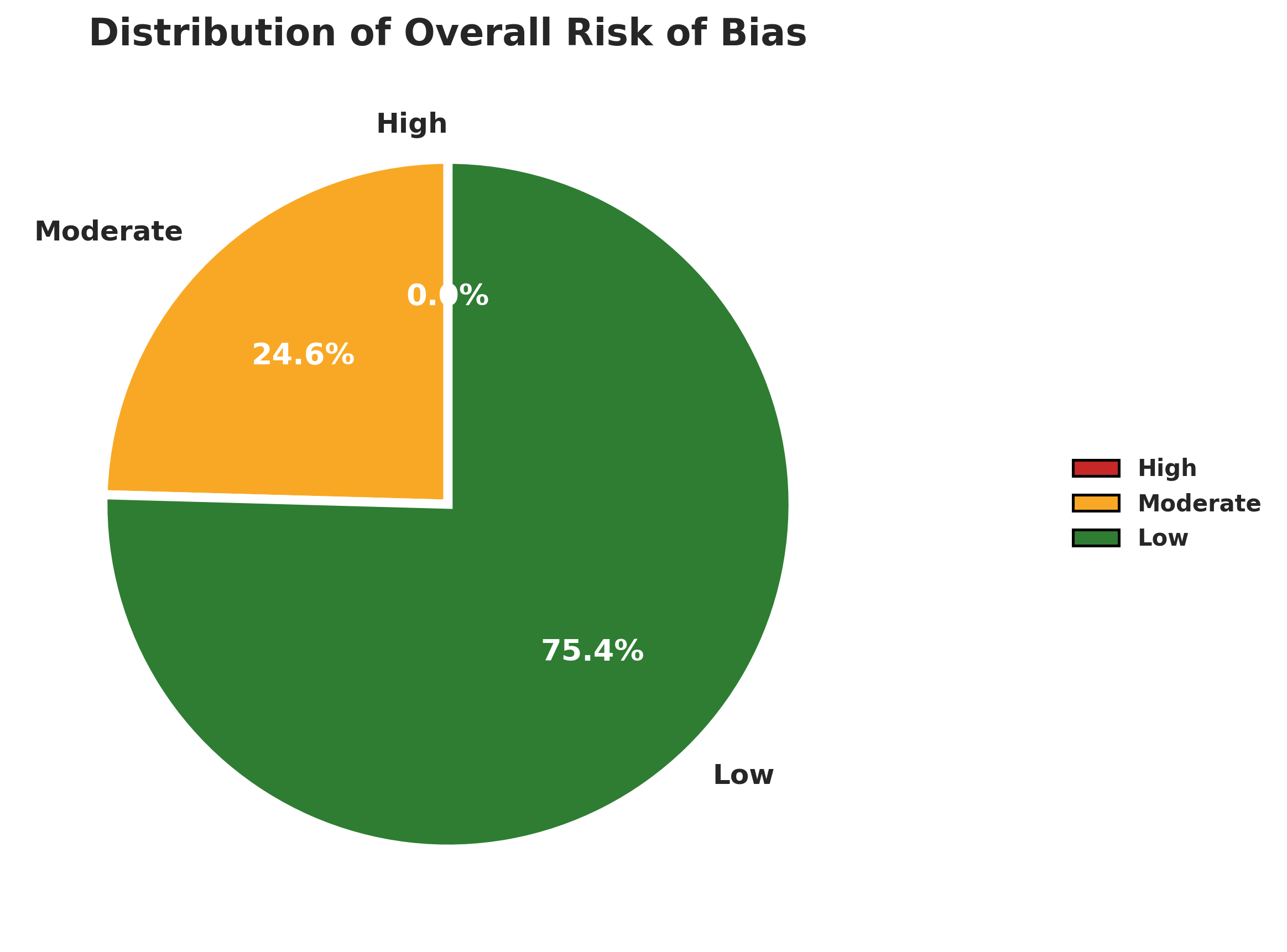

(B) Domain Heatmap of Included Studies

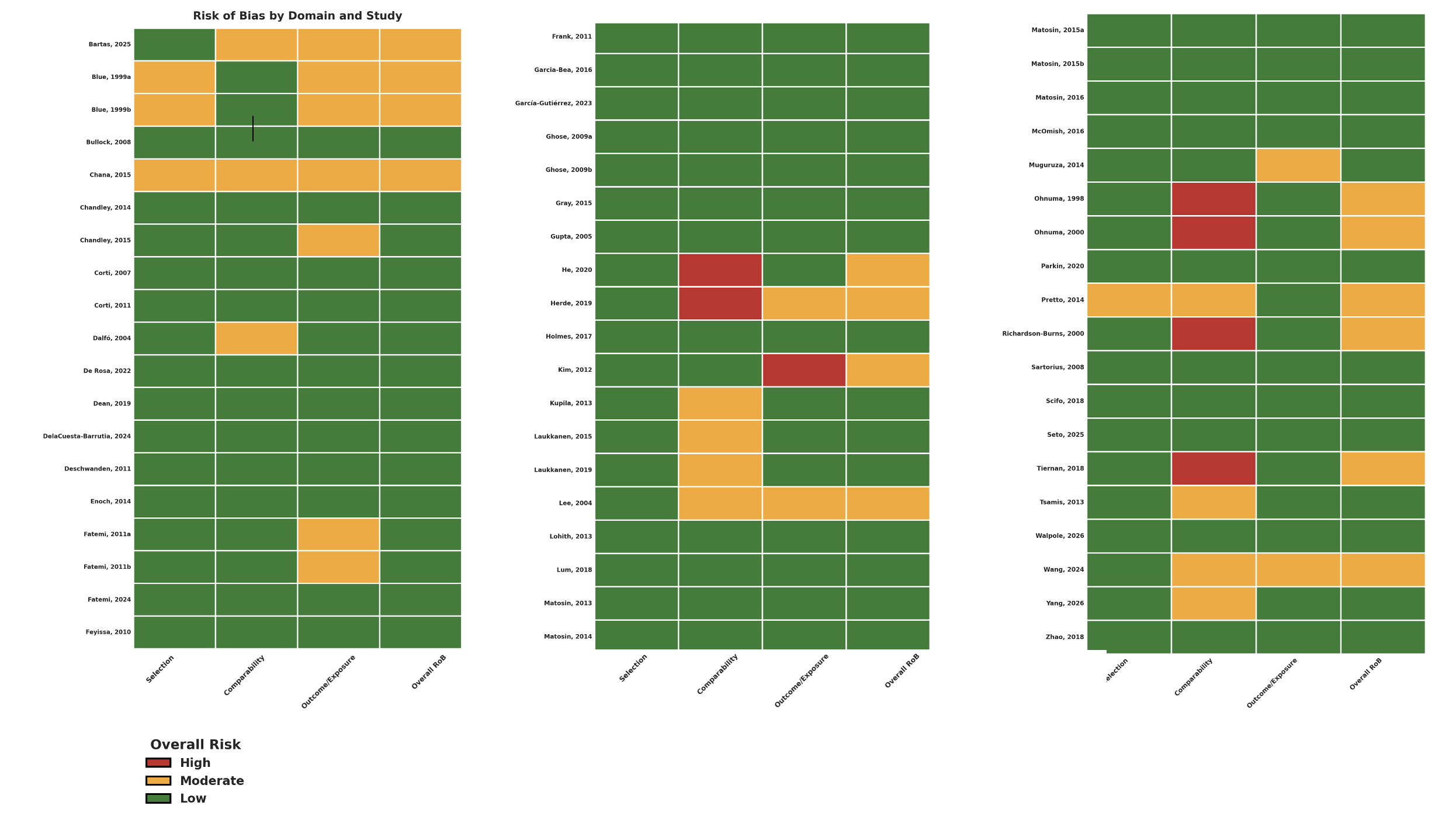

Abbreviation: Rob, Risk of bias.

**Supplemental Figure S2.	Evidence matrix of postmortem mGluR findings in (A) Bipolar disorder and (B) Patients who died by suicide.**

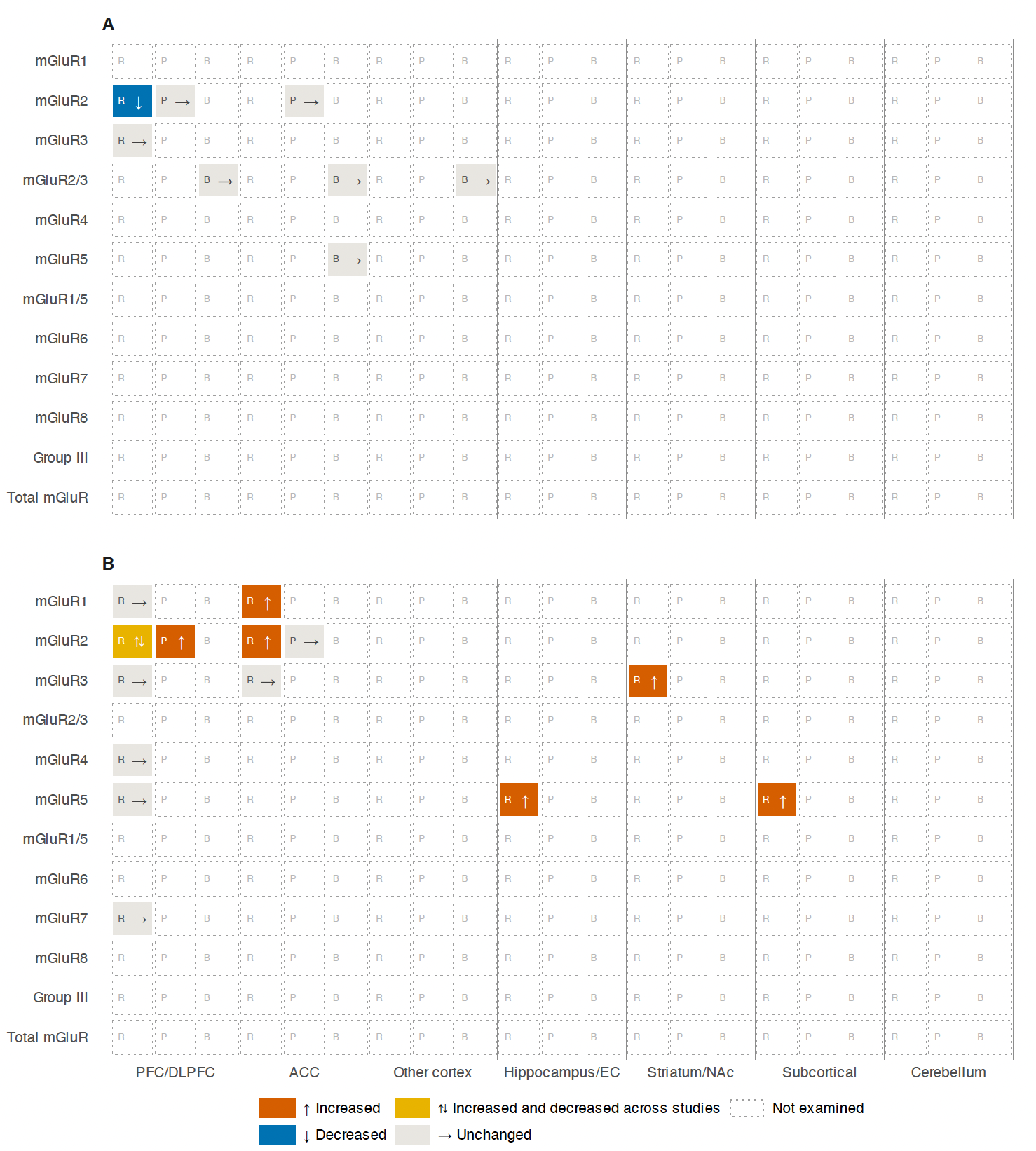

**Figure Legend:**
R, transcript abundance; P, protein abundance or immunoreactivity; B, radioligand binding.

Abbreviations: ACC: Anterior Cingulate Cortex; BA: Brodmann Area; DLPFC: Dorsolateral Prefrontal Cortex; EC: Entorhinal Cortex; mGuR: Metabotropic Glutamate Receptor; NAc: Nucleus Accumbens; PFC: Prefrontal Cortex.

**Supplemental Figure S3.	Evidence matrix of postmortem mGluR findings in (A) Alzheimer’s disease and (B) Diffuse Lewy Body Disease.**

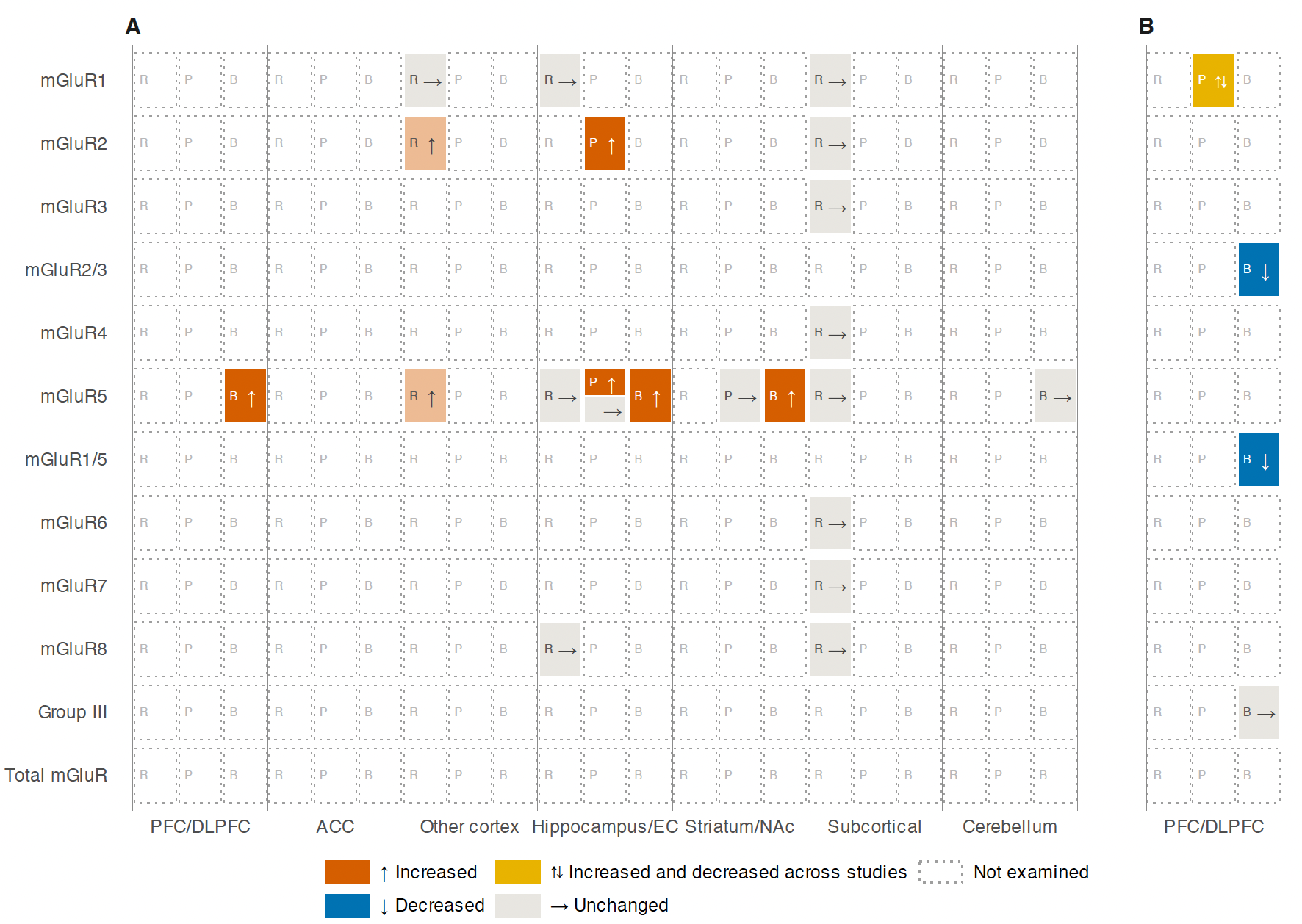

**Figure legends:**
R, transcript abundance; P, protein abundance or immunoreactivity; B, radioligand binding. A paler shade of any colour indicates a finding reported as nominal, or as not significant after correction for multiple comparisons.

Abbreviations: ACC: Anterior Cingulate Cortex; BA: Brodmann Area; DLPFC: Dorsolateral Prefrontal Cortex; EC: Entorhinal Cortex; mGuR: Metabotropic Glutamate Receptor; NAc: Nucleus Accumbens; PFC: Prefrontal Cortex.

**Supplemental Figure S4.	Evidence matrix of postmortem mGluR findings in (A) Fragile X-related conditions, (B) Rett syndrome, (C) Post-traumatic stress disorder.**

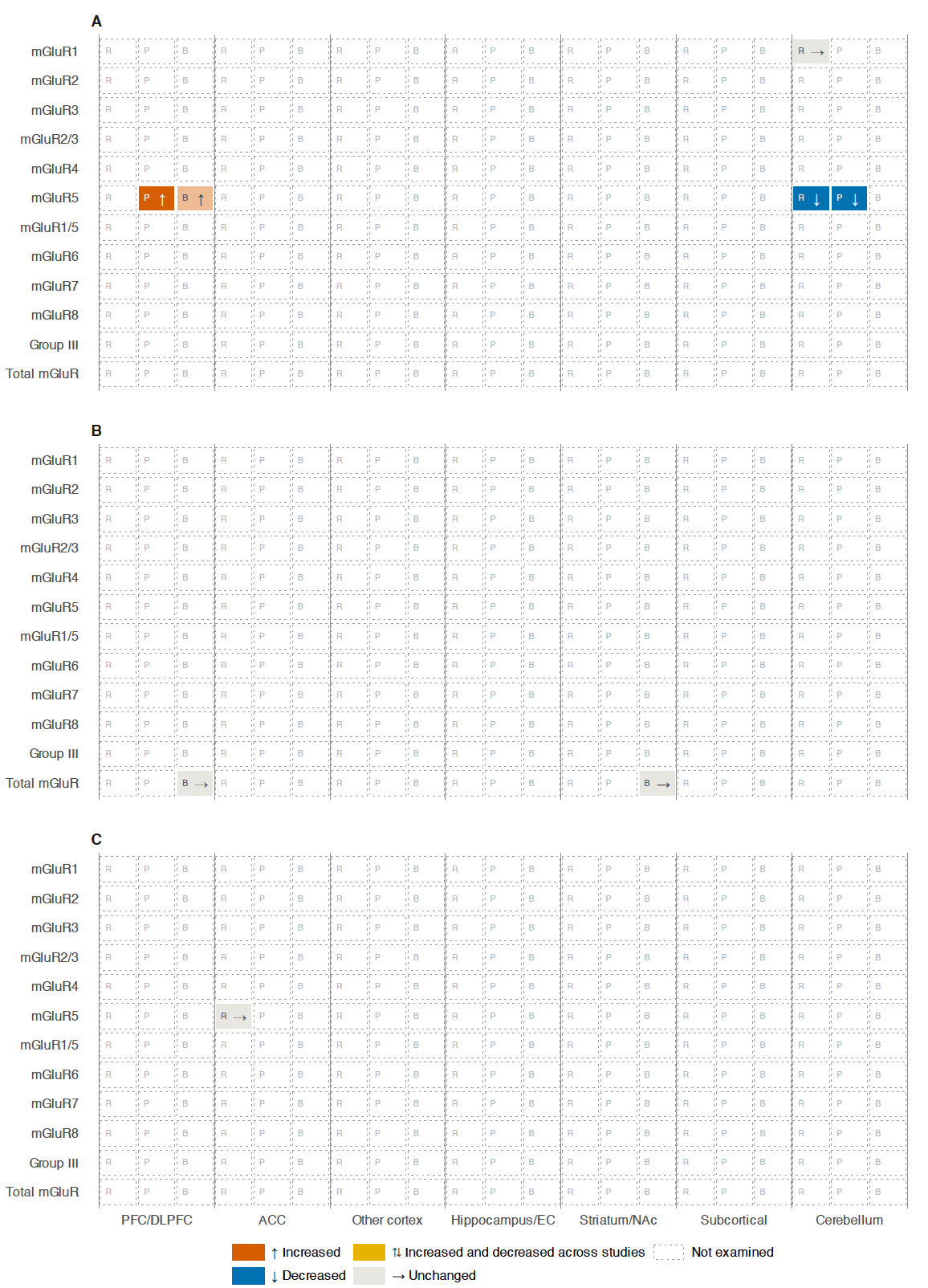

**Figure Legend:**
R, transcript abundance; P, protein abundance or immunoreactivity; B, radioligand binding. A paler shade of any colour indicates a finding reported as nominal, or as not significant after correction for multiple comparisons.

Abbreviations: ACC: Anterior Cingulate Cortex; BA: Brodmann Area; DLPFC: Dorsolateral Prefrontal Cortex; EC: Entorhinal Cortex; mGuR: Metabotropic Glutamate Receptor; NAc: Nucleus Accumbens; PFC: Prefrontal Cortex.

**Supplemental Figure S5.	Evidence matrix of postmortem mGluR findings in Autism spectrum disorder.**

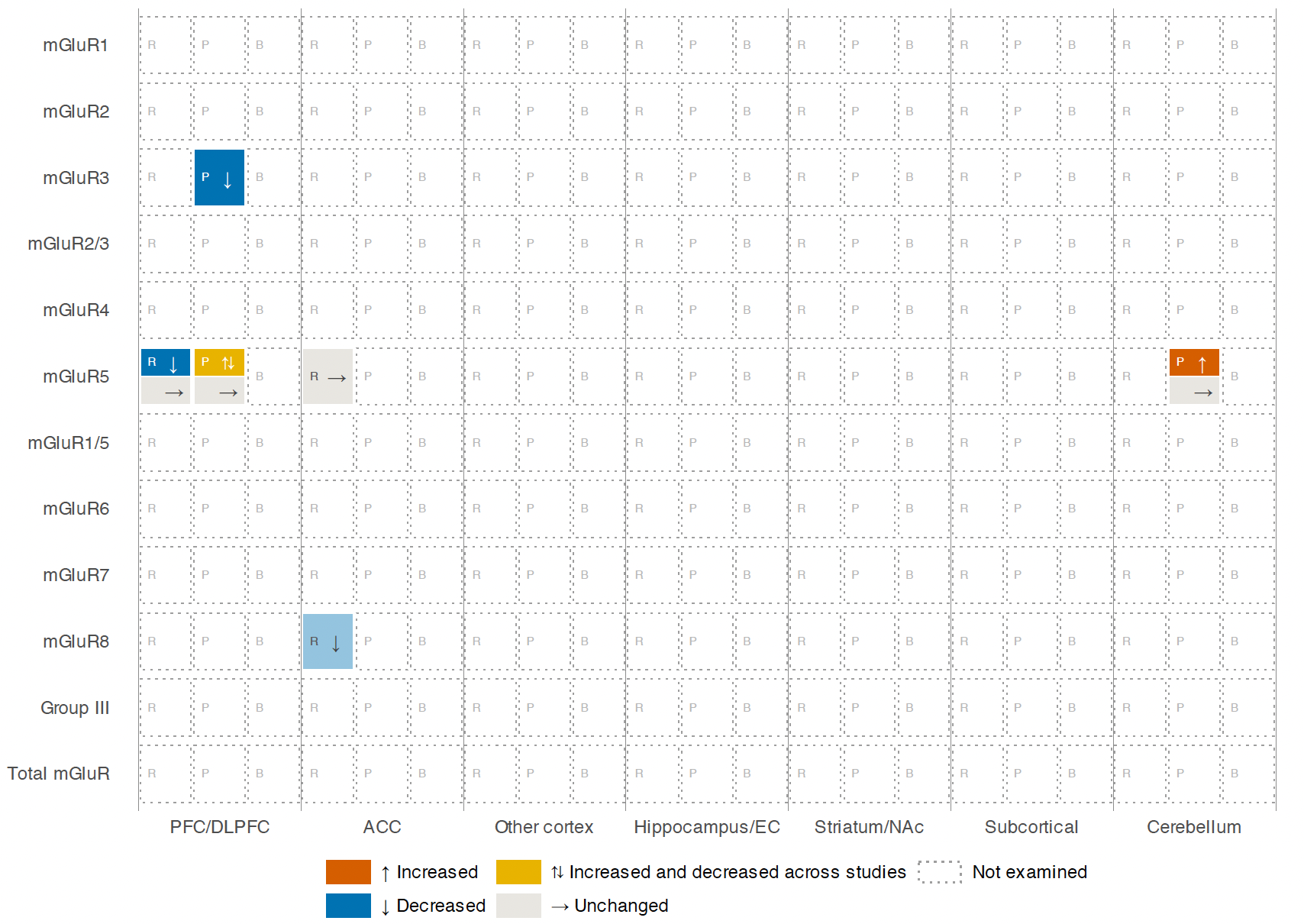

**Figure Legend:**

R, transcript abundance; P, protein abundance or immunoreactivity; B, radioligand binding. A paler shade of any colour indicates a finding reported as nominal, or as not significant after correction for multiple comparisons.

Abbreviations: ACC: Anterior Cingulate Cortex; BA: Brodmann Area; DLPFC: Dorsolateral Prefrontal Cortex; EC: Entorhinal Cortex; mGuR: Metabotropic Glutamate Receptor; NAc: Nucleus Accumbens; PFC: Prefrontal Cortex.

**Supplemental Figure S6.	Evidence matrix of postmortem mGluR findings in Alcohol use disorder.**

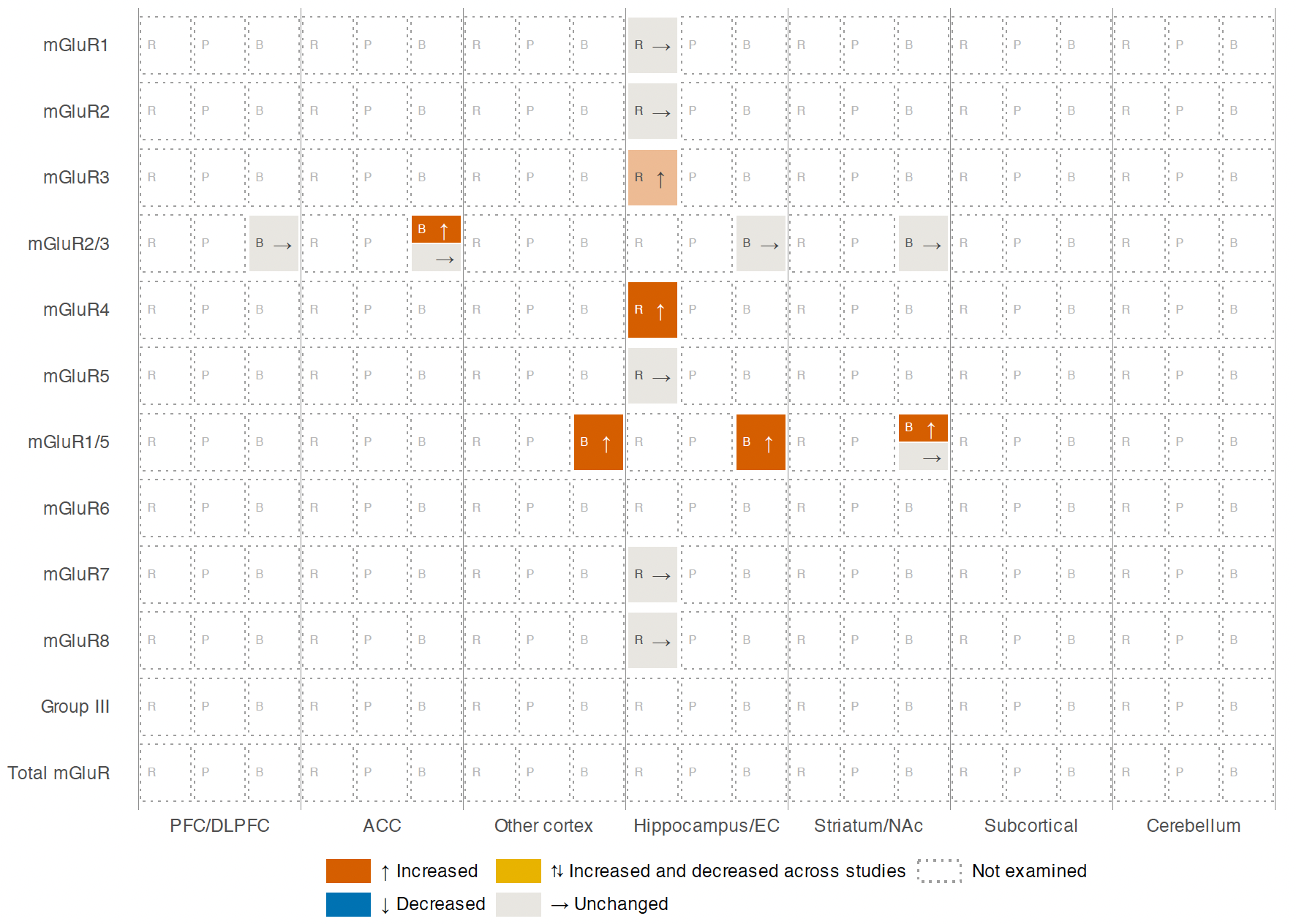

**Figure Legend:**

R, transcript abundance; P, protein abundance or immunoreactivity; B, radioligand binding. A paler shade of any colour indicates a finding reported as nominal, or as not significant after correction for multiple comparisons.

Abbreviations: ACC: Anterior Cingulate Cortex; BA: Brodmann Area; DLPFC: Dorsolateral Prefrontal Cortex; EC: Entorhinal Cortex; mGuR: Metabotropic Glutamate Receptor; NAc: Nucleus Accumbens; PFC: Prefrontal Cortex.
